# Connecting higher order interactions with ecological stability in experimental aquatic food webs

**DOI:** 10.1101/2023.05.04.539390

**Authors:** Chenyu Shen, Kimberley Lemmen, Jake Alexander, Frank Pennekamp

## Abstract

Community ecology is built on theories that represent the strength of interactions between species as pairwise links. Higher order interactions occur when the presence of a third (or more) species changes the pairwise interaction between a focal pair. Recent theoretical work has highlighted the stabilizing role of higher order interactions for large, simulated communities, yet it remains unclear how important higher order effects are in real communities. Here we used experimental communities of aquatic protists to examine the relationship between higher-order interactions and community stability (as measured by the persistence of species in a community). We cultured a focal pair of consumers in the presence of additional competitors and a predator and collected time series data of their abundances. We then fitted competition models with and without HOIs to measure interaction strength between the focal pair across different community compositions. We used survival analysis to measure the persistence of individual species. We found evidence that additional species positively affected persistence of the focal species and that HOIs were present in most of our communities. However, persistence was only linked to HOIs for one of the focal species. Our results vindicate community ecology theory positing that species interactions may deviate from assumptions of pairwise interactions, opening avenues to consider possible consequences for coexistence and community stability.

## Introduction

Community ecology seeks to unravel the mechanisms that maintain the tremendous organismal diversity on our planet. Species interactions, such as competition and predation, are important processes determining community structure and ecosystem stability and hence are important drivers of diversity (HilleRisLambers et al., 2012; Hooper et al., 2005; Landi et al., 2018). Most research on species coexistence has focused on pairwise interactions (Levine et al., 2017). However, in nature, species rarely interact only in a direct pairwise fashion, but rather in large networks of interacting species where a suite of indirect effects are likely to be important (Levine et al., 2017). Recently, these indirect effects have gained increasing attention, since in some cases they may explain the dynamics and stability of communities better than classic pairwise models (Grilli et al., 2017; Letten & Stouffer, 2019; Mayfield & Stouffer, 2017).

Lotka-Volterra competition or predator-prey models assume that interaction is an intrinsic property of two species interacting in a given environment, and therefore the parameters that determine the strength of interactions are independent of the community in which these species are embedded (Werner & Peacor, 2003). However, when more than two species co-occur (Worthen & Moore, 1991), indirect effects can arise through chains of pairwise interactions, or through higher-order interactions, that is changes of the per capita effect of one competitor on another in the presence of additional species (Levine et al., 2017; Wootton, 1994). Since interaction chains and higher-order interactions emerge from fundamentally different mechanisms, it is important to understand their underlying mechanisms for appropriate modelling and inference (Levine et al., 2017).

Interaction chains emerge when pairwise interactions are embedded in a network of interactions. A series of such direct interactions between species pairs can lead to connections between species that do not directly interact with each other (Wootton, 1993). Despite introducing indirect pathways between species, interaction chains are the result of a series of fixed, strictly pairwise interactions. In contrast, “higher-order interactions” (HOIs (Case & Bender, 1981; Letten & Stouffer, 2019; Mayfield & Stouffer, 2017); also called “interaction modifications” (Abrams, 1983)), encompass the non-additive effects that interactions between individuals of different, or the same species have on the per capita effect of one species on the focal species. Wootton (1994) observed both types of indirect effects in the upper zone of a rocky intertidal community. As an example of an interaction chain, bird predators indirectly increase acorn barnacle (*Balanus glandula*) abundance by consuming limpets (*Lottia digitalis*), which dislodge or consume young acorn barnacles. An example of an interaction modification, or inter-specific HOI, is barnacles altering the bird-limpet interaction by changing the ability of birds to find limpets due to the similar color of *L. digitalis* and barnacles shells. HOIs are most commonly described in terms of how a third species impacts the interaction between a focal species pair. However, intraspecific HOIs are also important to consider and represent the cumulative impacts of interactions among the individuals of each ‘competitor species’ (that is, intraspecific crowding) on the focal species (Letten & Stouffer, 2019; Mayfield & Stouffer, 2017). In the intertidal community, intraspecific HOIs could occur when birds influence the foraging behavior of barnacles, causing them to aggregate in areas sheltered from bird predation, which leads to increased intraspecific competition. Interaction chains can be predicted with only a knowledge of pairwise species interactions; in contrast, descriptions of intra- and interspecific HOIs require a knowledge of all species combinations involved (Wootton, 1994).

Theoretical work has suggested that empirical studies of interactions between species should consider HOIs (Grilli et al., 2017) given that simple models of pairwise interactions fail to explain the stable persistence in simulation models of very large ecological communities (Barabás et al., 2016; Clark, 2010; Gibbs et al., 2022; Kleinhesselink et al., 2022; Levine et al., 2017). Recently, many theoretical advances have been made to understand the influence of HOIs on community dynamics (Gibbs et al., 2022; Grilli et al., 2017; Letten & Stouffer, 2019; Mayfield & Stouffer, 2017; Singh & Baruah, 2021). A particular focus has been on the effect of HOIs on the stability of communities, specifically the persistence of all species in a community. However, simulation-based studies often consider highly complex and diverse communities (Bairey et al., 2016; Singh & Baruah, 2021), making it impractical to verify their findings through experimental manipulations.

A valuable approach to simplify the complexity of communities is to analyze their component community modules (Holt, 1997). Studies of community modules have revealed how interactions between species affect species persistence and community stability (Kondoh, 2008; Mayfield & Stouffer, 2017; McCann et al., 1998). The classical ecological theory holds that strong consumer–resource interactions can lead to destabilization and potential collapse of the community due to overexploitation (Rosenzweig, 1971). Theoretical investigations on food web modules have highlighted the crucial role of weak interactions, especially omnivorous links, in maintaining community stability (Emmerson & Yearsley, 2004; McCann et al., 1998). Experimental manipulations of these modules have confirmed that weak interactions, coupled with strong interactions mediated by generalist consumers, enhanced community stability by reducing interaction strength (Rip et al., 2010). A recent study on community modules of the zebrafish gut microbiome has also confirmed that HOIs weaken the intense pairwise competition between bacteria, thereby stabilizing the community and promoting coexistence (Sundarraman et al., 2020).

Despite the theoretical support for the importance of higher order interactions for coexistence and community stability, empirical evidence connecting HOIs with ecological stability remains very scarce (but see Mayfield & Stouffer, 2017; Mickalide & Kuehn, 2019; Sundarraman et al., 2020)). This is partly due to the challenges of empirically quantifying HOIs (Billick & Case, 1994), which requires the evaluation of the interaction strength between two species accounting for changes in density or the presence of additional species (Adler et al., 2018). Even in simple communities, this can quickly become logistically infeasible. In addition, the quantification of the stability of communities requires time-series to evaluate species persistence over extended periods of time, presenting a second logistical hurdle.

Our goal was to experimentally test the causal links between higher-order interactions and the persistence of species within communities. Microcosms are a convenient tool to investigate concepts in community ecology because they are easily manipulated, can be highly replicated, and have large population sizes (Altermatt et al., 2015). Moreover, a range of analytical approaches is available to estimate intra- and interspecific interaction strengths within these systems (Carrara et al., 2015). We, therefore, used microcosms to experimentally examine HOIs and community persistence in aquatic microbial communities. We used six different community compositions, which all included a focal pair of ciliate consumers competing for the same resource. We tested the effect of two additional competitors as well as the presence of a predator on the interaction strength between the focal pair. Time series of population dynamics were collected to quantify the strength of species interactions and examine the effect of HOIs on community stability and address the following questions:

1. Are higher order interactions detectable in our communities?
2. Do HOIs differ depending on the identity and trophic role of the third species?
3. Is there a relationship between the presence of HOIs and community persistence?

We hypothesized that additional competitors would influence the effect of the two focal species on each other since all species engage in resource competition. For instance, in the presence of a third species, focal species may shift their foraging behaviour and thereby their pairwise effect on one another. We also expected that higher-order effects would arise in communities with a predator whose behaviour is affected by different consumer body sizes which could render one prey susceptible to predation while the other may be attacked but is too large to be consumed. All else being equal, we expected a stabilizing effect on the persistence of the populations of the two focal species for HOIs that decreased their interaction strength.

## Methods

### Data collection: Community Experiment

#### Food web construction and culture conditions

To experimentally test higher order interactions, we constructed an experimental food web. The ciliates of our microbial food web were primarily bacterivorous (Pennekamp et al., 2018), thus the experimental communities were sustained on a mix of three species of bacteria (*Bacillus subtilis, Serratia fonticola and Brevibacillus brevis*) decomposing the protozoan pellet medium (PPM; provided by Carolina Biological Supplies, Burlington NC, USA; concentration of 0.55 g/L, see Altermatt et al., 2015).

Four bacterivorous ciliates (*Colpidium striatum, Dexiostoma campylum, Paramecium caudatum* and *Spirostomum teres*) constitute the intermediate consumers in our experimental microcosms. Both *Colpidium striatum* (50-100 um) and *Dexiostoma campylum* (35-90 um) are consumed by the top predator *Spathidium sp.* (40 - 300 um), which cannot survive only on bacteria as prey (Woodruff & Spencer, 1921). Due to the large body size of *Paramecium caudatum* (170–300 um) (Foissner & Berger, 1996) and *Spirostomum teres* (150–400 um) (Bick & World Health Organization, 1972), we expected these species to interfere with the predator because they are more difficult or impossible to consume; however, depending on the mode of attack of the predator, even failed consumption could result in death of prey.

We chose these five eukaryotic ciliates (hereafter referred to by the genus name only) because their pairwise trophic interactions are well documented by previous studies (Daugaard et al., 2019; Pennekamp et al., 2018; Tabi et al., 2019). The consumptive and competitive interactions present in our experimental food web are shown in Figure 1.

**Figure 1:**
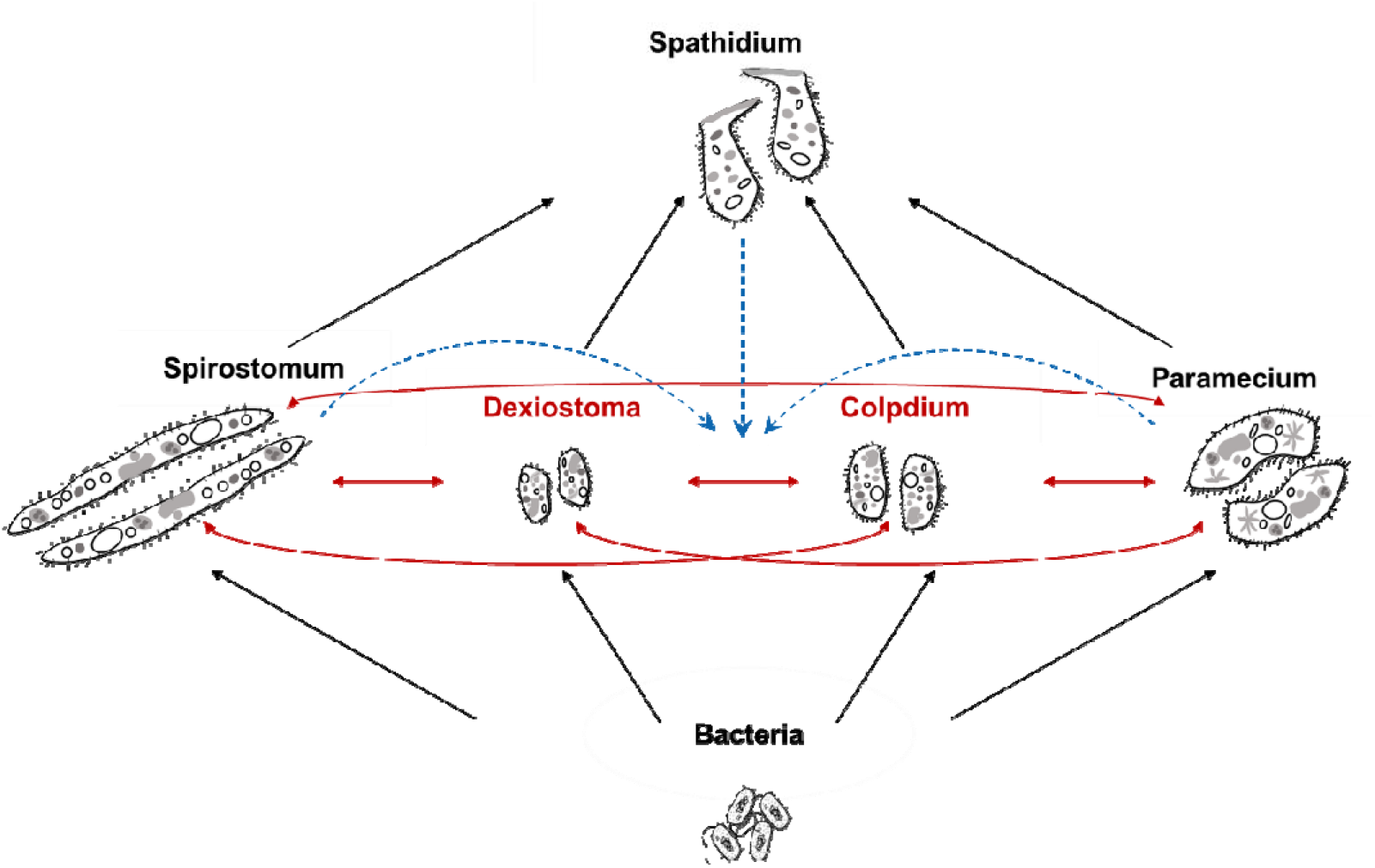
Microbial food web in our experiment. Consumptive interactions (in black) among the top predator (*Spathidium sp.*), the consumers *(Colpidium striatum, Dexiostoma campylum*, *Paramecium caudatum* and *Spirostomum teres*) and a mix of three bacteria species (*Bacillus subtilis, Serratia fonticola and Brevibacillus brevis*). All the consumers compete with each other (red arrows), with the focal pair marked in red. Hypothesized HOIs are indicated by the blue dashed lines with arrows.

#### Experimental design

To assess how the interaction strength between two focal species is altered by adding a third or more species, we established the following treatments: (1) two prey (*Colpidium* and *Dexiostoma*) alone to investigate the direct effects between the two focal species (CD) (2) two prey and *Paramecium* (CDP). (3) two prey and *Spirostomum* (CDS). These communities were also grown in the presence of the top predator *Spathidium (*CDPd, CDPPd, CDSPd; predator = Pd) (Figure 2), yielding six treatments in total. For each treatment, we cultured four replicates resulting in 24 microcosms in total.

**Figure 2:**
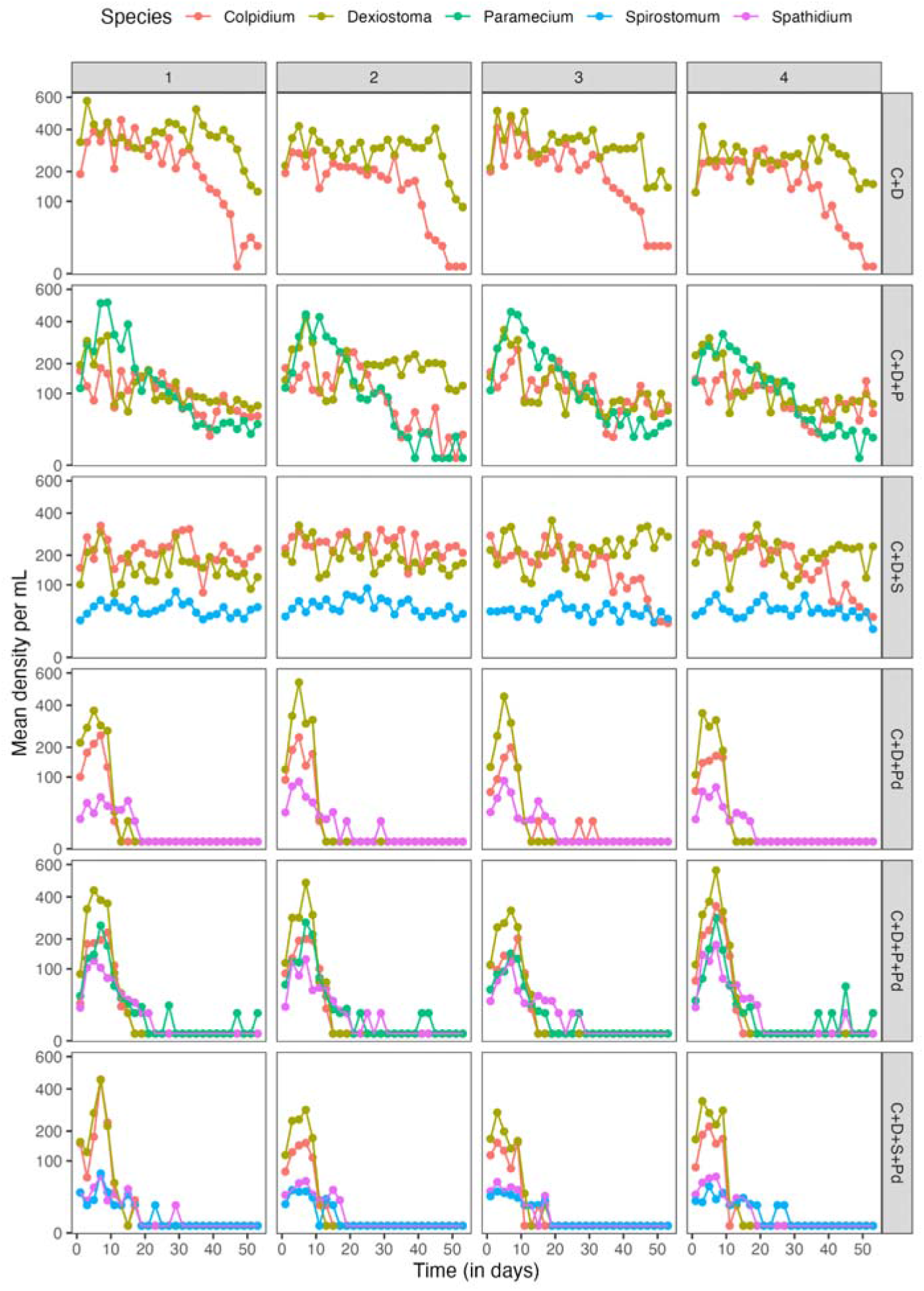
Population dynamics for each experimental community. Rows are six different treatments in our experiment. Each treatment is represented by the species it contains (C for *Colpidium*, D for *Dexiostoma*, P for *Paramecium*, S for *Spirostomum* and Pd for predator). The y-axes are plotted on a logarithmic scale. Results for each replicate are shown in columns. Most species in competition treatments (first three rows) survived to the last day of measurement but rapidly dropped below detection limits when housed with predators (last three rows).

To start the experiment, we took out 20 or 30 mL of the bacterized PPM medium (depending on experimental community composition) and replaced it with 10 mL of the stock culture of each consumer species at their carrying capacity. Therefore, the starting densities of all consumers were 10% of their carrying capacities. In the predation treatments, four days after the introduction of the consumer species ten predator individuals were pipetted from the maintenance plates to the microcosms. To assure establishment in each microcosm, another ten individuals of predators were added after 8 days. The experiments were conducted at 15°C with no light, and suitable growing conditions for ciliates (Altermatt et al., 2015).

#### Video sampling and species classification

Experimental units were sampled every second day for 53 days to capture time series of the dynamical changes in the abundance of ciliates. For each sampling event, the microcosm was gently agitated, a subsample of 250μL was mounted onto a glass slide and covered with a glass lid. Three five-second videos (at 25 frames per second) were taken using 25× magnification on a stereomicroscope (Leica M205 C) mounted with a digital CMOS camera (Hamamatsu Orca Flash 4.0 C11440, Hamamatsu Photonics, Japan) with dark field illumination. We took three subsamples for each microcosm to get a precise estimate of abundance and processed the videos with the R package BEMOVI (version 1.0.2) (Pennekamp et al., 2015). We took the mean of the three videos as our measure of density. If no individuals were detected, we assigned a zero. The volume lost from the microcosm by sampling during the course of the experiment was replaced with a fresh bacterized PPM medium.

For species classification, we trained a support vector machine classifier (SVM) on 350 to 400 randomly chosen and manually labelled trajectories of each species across the community compositions and time with the R package e1071 (Meyer et al., 2022). We also included a “noise” class in our classifier representing spurious trajectories due to background movement. Twenty morphological and movement features extracted from an established classification pipeline (Pennekamp et al., 2017) were selected to train the support vector machine to distinguish among classes based on information about body size and movement patterns (Table S1). As ciliate phenotypes may change over time (Pennekamp et al., 2017), we included “week number” to enhance the accuracy of the classifier. Further details about the classification can be found in the supplementary material.

### Data analyses: interaction strengths and community stability

#### Abundance

To understand the effect of community composition on the abundance of the species, we calculated the mean abundance over the entire 53-day experiment for each species in each of the six communities. Mean abundance of each of the five species was analysed with two-way ANOVA (predictors: identity of the third species [levels: *Paramecium* or *Spirostomum*] and the presence of the predator [levels: present or absent]).

#### Estimating species interaction strengths

We estimated the effect of species j on the population growth rate of species i within a regression framework, where the population growth rate of species i is regressed against the densities of species i and j to determine the intra- and interspecific effects on the population growth rate of species i (Pfister, 1995). The *per capita* population growth rate of each species i was calculated as 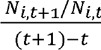 where N_i,_(t+l)-t_t+1_ is the abundance at time t+1 and N_i,t_ is abundance at time t. We used the gauseR package to calculate the per capita population growth rate (Mühlbauer et al., 2020). To explore how interaction strength and HOIs may differ between communities, we compared the fit of six models to the per capita population growth rate of the focal species (either *Dexiostoma* or *Colpidium*). First, we fitted the Lotka-Volterra model with no HOIs:

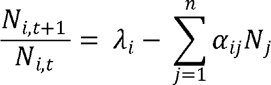

For the simple additive model, λ_i_ is the intrinsic population growth rate and α_ij_ represents both the intra- and interspecific interaction coefficients. To test for intra- and interspecific HOIs, we then added interaction terms to the additive model LV, resulting in the interactive LV model (Letten & Stouffer, 2019). For the interactive LV model only including intra-specific HOIs, we included the interaction term β_ijj_ for the effect of species j on i, where β_ijj_ captures the cumulative impacts of intra-specific interactions on the focal species:

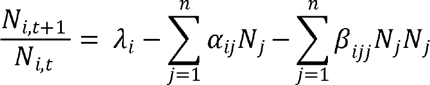

The identity of species j may or may not be the same as species i. Next, we fitted an interactive LV model that only includes interspecific HOIs, where β_ijk_ captures the cumulative impacts of interspecific interactions on the focal species:

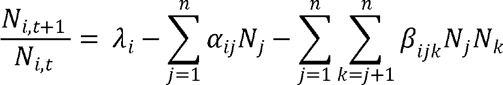

Here, the identity of species k strictly excludes the focal species i.

Finally, we tested the fit of the interactive LV model including both intra- and interspecific HOIs (full HOI model):

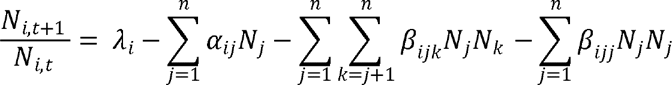

To test whether the shape of the density-dependence deviates from the linear form of the additive Lotka-Volterra model, we fitted the Ricker model with nonlinear density dependence. First, we fitted the additive model without HOIs:

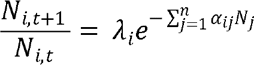

and interactive form only including interspecific HOIs (terms as previously defined):

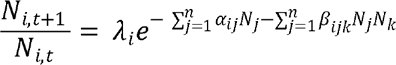

We used generalized linear models with the “identity” link function assuming Gaussian errors to estimate the coefficients of the additive and interactive LV model, considering all errors as measurement errors. For the Ricker model, we used a GLM with the “log” link function, assuming also Gaussian errors. For the fitted models, the intercept can be interpreted as the maximum intrinsic population growth rate, while the regression coefficients represent the *per capita* effect of species i on itself and the per capita effects of species j on species i. Interaction terms describe the mediating effects that the density of species j can have on the effect of species i (and vice versa) on the population growth rate of the focal species (Letten & Stouffer, 2019). We performed an AICc model selection adjusting for the small sample size when identifying the model that best explained the population growth rate of each focal species (Burnham & Anderson, 2002). AICc was preferred over the Bayesian information criterion since the models which we fit are approximate and we do not necessarily expect to include the true model in our set (Aho et al., 2014). We then calculated the difference (ΔAICc) between the model with the lowest AICc and all other models. Models with a ΔAICc larger than 2 are considered to be different, thus the model with the fewest parameters and a ΔAICc less than 2 is considered the most parsimonious model. Since the presence of predators drove some prey to early extinction, we only estimated interaction strengths in communities without predators.

#### Estimating species survival (persistence)

We measured the persistence of the focal species *Colpidium* and *Dexiostoma*, as well as the predator *Spathidium* as a proxy of the population stability of these species using non-parametric survival analysis. We estimated the time to event (i.e., extinction), accounting for right-censoring, that is, the possibility of missing future extinctions since we only recorded the abundance for 53 days. We tested the effect of community composition on the Kaplan-Meier estimate of the focal species with log-rank tests, followed by pairwise tests of community composition in case of a significant community wide effect with the survminer package (Kassambara et al., 2021).

All of the above analyses including the species classification were performed with the statistical computing environment R (R Core Team, 2022).

## Results

### Population dynamics over time

The population dynamics of the protists differed depending on the community context (Figure 2). When *Colpidium* and *Dexiostoma* competed only with each other, we consistently observed the extinction of *Colpidium* towards the end of the experiment, while *Dexiostoma* persisted. The addition of *Paramecium* or *Spirostomum* led to a longer period of coexistence of *Dexiostoma* and *Colpidium*. The addition of the top predator *Spathidium* considerably shortened the persistence of all species. Thus, the addition of a third competitor increased the period of coexistence of the focal species pair, whereas the addition of the predator led to the rapid collapse of the whole community.

The addition of the competitor *Paramecium* had a negative effect on the mean abundance of both *Colpidium* (b=-0.60, 95% CI = −0.89 to −0.32, n=4) and *Dexiostoma* (b= −0.81, 95% CI = −1.0 to −0.59, n=4, figure 3). In contrast, adding the competitor *Spirostomum* only had a negative effect on the mean abundance of *Dexistoma* (b= −0.43, 95% CI = −0.67 to −0.19, n=4, figure 3).

**Figure 3:**
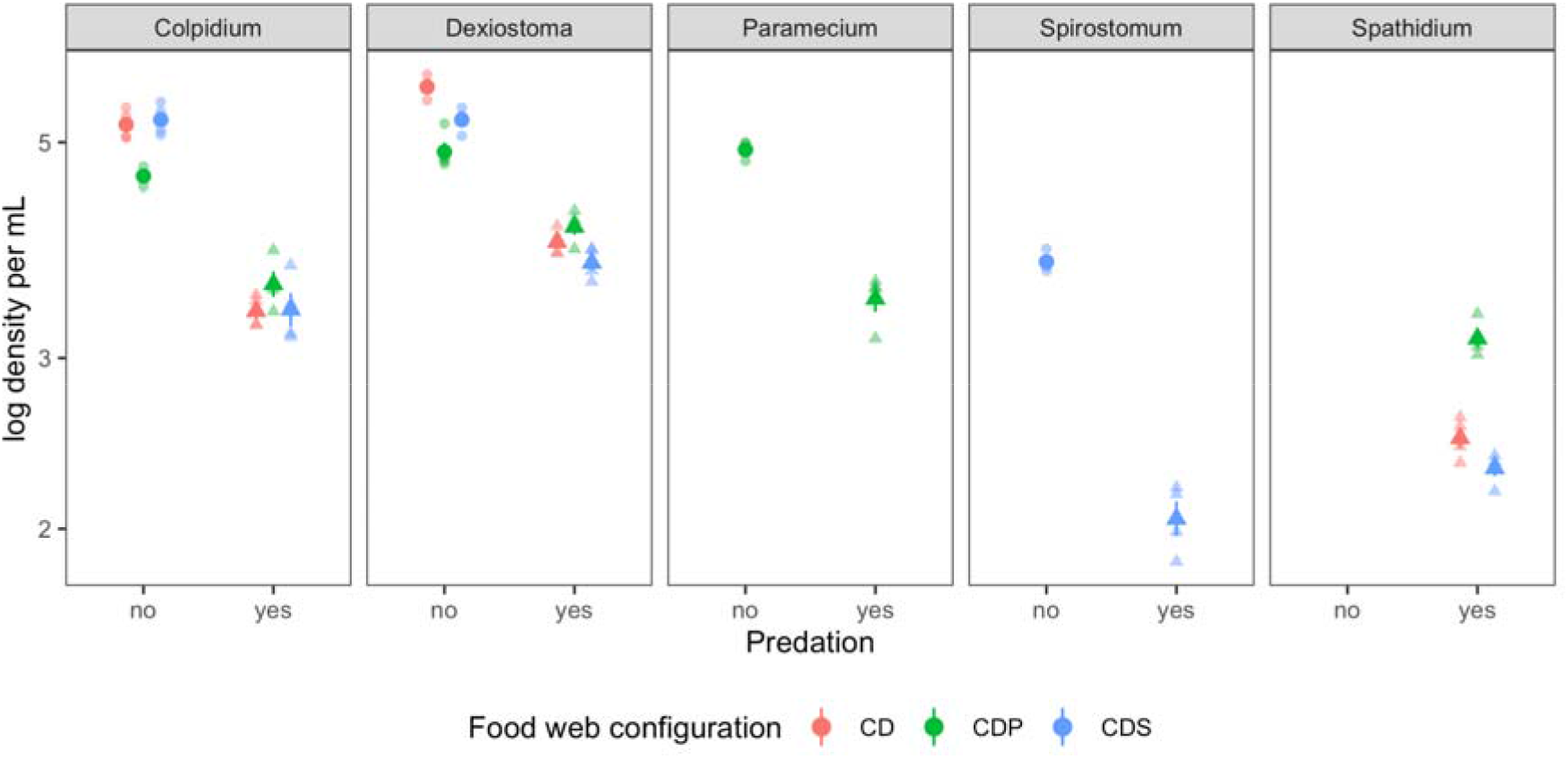
Density of focal species as a function of community composition. Each panel shows one of the protist species. Whether the predator was present is shown on the x-axis. The treatments with predators are shown with triangles, and the treatments without predators are shown with circles. The three competitive communities are shown in color (CD = *Colpidium* & *Dexiostoma*, CDP = *Colpidium*, *Dexiostoma* & *Paramecium*, CDS = *Colpidium*, *Dexiostoma* & *Spirostomum*). The solid shapes show the mean and the error bars show the standard error of the mean (calculated across the duration of the experiment), and the transparent shapes show the actual data points.

The addition of the predator had a negative effect on the mean abundance of all competing protist species (figure 3). In communities composed of just the focal pair the presence of the predator resulted in a similar reduction in mean abundance for both *Dexiostoma*: (b= −1.8, 95% CI = −2.0 to −1.5, n=4) and *Colpidium* (b= −1.9, 95% CI = −2.1 to −1.6, n=4, figure 3). In communities with a third competitor, the predator had a greater negative effect on the mean abundance of *Spirostomum* (b= - 1.7, 95% CI = −1.9 to −1.5, n=4) than *Paramecium* (b= −1.5, 95% CI = −1.7 to −1.2, n=4, figure 3). The addition of *Spirostomum* had no effect on the mean abundances of *Dexiostoma*, *Colpidium* and the predator compared to the focal pair cultured with just the predator. Intriguingly, the presence of both *Paramecium* and the predator had a positive effect on the density of both *Colpidium* (b= 0.83, 95% CI = 0.43 to 1.2, n=4) and *Dexiostoma* (b= 0.96, 95% CI = 0.62 to 1.3, n=4, figure 3) in comparison to when the focal species were exposed to the predator in isolation. This translated into higher density for the predator *Spathidium* when *Paramecium*, *Colpidium* and *Dexiostoma* were present (b= 0.66, 95% CI = 0.48 to 0.84, n=4, figure 3).

### Higher-order interactions

Model selection revealed that higher order interactions influenced the *per capita* population growth rates of *Dexiostoma* and *Colpidium* in some communities (Table 1, Figures S1-7). For *Colpidium*, the interactive LV model only including intraspecific HOIs was best supported in the two-species community of *Colpidium* and *Dexiostoma*, indicating that *Dexiostoma* modified the effect of *Colpidium* on itself. In culture with a third species, the best supported model differed: when *Colpidium* was present with *Dexiostoma* and *Paramecium*, the additive Ricker model was the most parsimonious (ΔAIC = 0.06 but fewer parameters). In contrast, when *Colpidium* was present with *Dexiostoma* and *Spirostomum*, the interactive Ricker model was best supported, suggesting interspecific HOIs. For the *per capita* population growth rate of *Dexiostoma*, the interactive LV model only including intraspecific HOIs (ΔAIC = 0.59 but fewer parameters) was most supported when only *Colpidium* was present, suggesting that *Colpidium* modified the effect of *Dexiostoma* on itself. When a third species was present, in both cases the additive Ricker model was best supported, suggesting that neither intra-nor interspecific HOIs were operating. Since all the best supported models were either the interactive LV or Ricker models, all intra- and interspecific interactions were nonlinear.

**Table 1:** Model selection table showing the AICc values for the per capita population growth rate of the two focal species across the three community compositions. Additive and interactive versions of the Lotka-Volterra (LV) and Ricker models were fitted to the data. K is the number of estimated parameters, ΔAICc the difference between the model with the lowest AICc and the AICc of the specific model.

| species | community | model | K | AICc | $\Delta AICc$ |
| --- | --- | --- | --- | --- | --- |
| Colp | CD | **interactive LV (intra HOI)** | **6** | **-48.74** | **0.00** |
|  |  | **interactive LV (full HOI)** | **7** | **-48.42** | **0.32** |
|  |  | additive Ricker | 4 | -41.81 | 6.93 |
|  |  | additive LV | 4 | -41.61 | 7.13 |
|  |  | interactive Ricker | 5 | -39.59 | 9.15 |
|  |  | interactive LV (inter HOI) | 5 | -39.39 | 9.35 |
|  | CDP | **interactive LV (intra HOI)** | **8** | **63.30** | **0.00** |
|  |  | **additive Ricker** | **5** | **63.37** | **0.06** |
|  |  | interactive Ricker | 8 | 68.21 | 4.90 |
|  |  | interactive LV (full HOI) | 11 | 68.81 | 5.51 |
|  |  | additive LV | 5 | 75.50 | 12.20 |
|  |  | interactive LV (inter HOI) | 8 | 77.96 | 14.65 |
|  | CDS | **interactive Ricker** | **8** | **-75.94** | **0.00** |
|  |  | **interactive LV (inter HOI)** | **8** | **-74.09** | **1.85** |
|  |  | interactive LV (full HOI) | 11 | -68.82 | 7.12 |
|  |  | additive Ricker | 5 | -66.40 | 9.54 |
|  |  | additive LV | 5 | -65.64 | 10.30 |
|  |  | interactive LV (intra HOI) | 8 | -61.48 | 14.46 |
| Dexio | CD | **interactive LV (full HOI)** | **7** | **-87.01** | **0.00** |
|  |  | **interactive LV (intra HOI)** | **6** | **-86.42** | **0.59** |
|  |  | additive Ricker | 4 | -80.56 | 6.46 |
|  |  | interactive Ricker | 5 | -79.61 | 7.40 |
|  |  | additive LV | 4 | -75.10 | 11.92 |
|  |  | interactive LV (inter HOI) | 5 | -73.87 | 13.14 |
|  | CDP | **additive Ricker** | **5** | **3.88** | **0.00** |
|  |  | additive LV | 5 | 5.96 | 2.08 |
|  |  | interactive LV (intra HOI) | 8 | 7.80 | 3.92 |
|  |  | interactive Ricker | 8 | 8.80 | 4.92 |
|  |  | interactive LV (inter HOI) | 8 | 10.12 | 6.25 |
|  |  | interactive LV (full HOI) | 11 | 12.03 | 8.16 |
|  | CDS | **additive Ricker** | **5** | **-60.05** | **0.00** |
|  |  | additive LV | 5 | -56.38 | 3.67 |
|  |  | interactive LV (intra HOI) | 8 | -56.21 | 3.84 |
|  |  | interactive Ricker | 8 | -54.32 | 5.73 |
|  |  | interactive LV (full HOI) | 11 | -51.22 | 8.83 |
|  |  | interactive LV (inter HOI) | 8 | -50.82 | 9.23 |

### Population persistence

In the absence of predators, *Colpidium* survival was contingent on the community composition (log-rank test p= 0.0066, n=4, figure 4A). Without predators, adding *Spirostomum* increased *Colpidium* survival (pairwise log-rank test p = 0.02, n=4, figure 4A), whereas adding *Paramecium* did not. In contrast, *Dexiostoma* persisted in the absence of predators regardless of the community composition (log-rank test p = 0.37, n=4, figure 4C). In the presence of predators, *Colpidium* survival did not depend on the identity of the competitor species (figure 4B). In contrast for *Dexiostoma*, the community composition did impact survival (log-rank test p = 0.024, n=4), since adding *Paramecium* to the community increased *Dexiostoma* survival (figure 4D). *Spathidium* survival depended on the prey community composition (log-rank test p = 0.011, n=4), with *Paramecium* increasing the survival of *Spathidium* (pairwise log-rank test p = 0.018, n=4, figure 4E).

**Figure 4:**
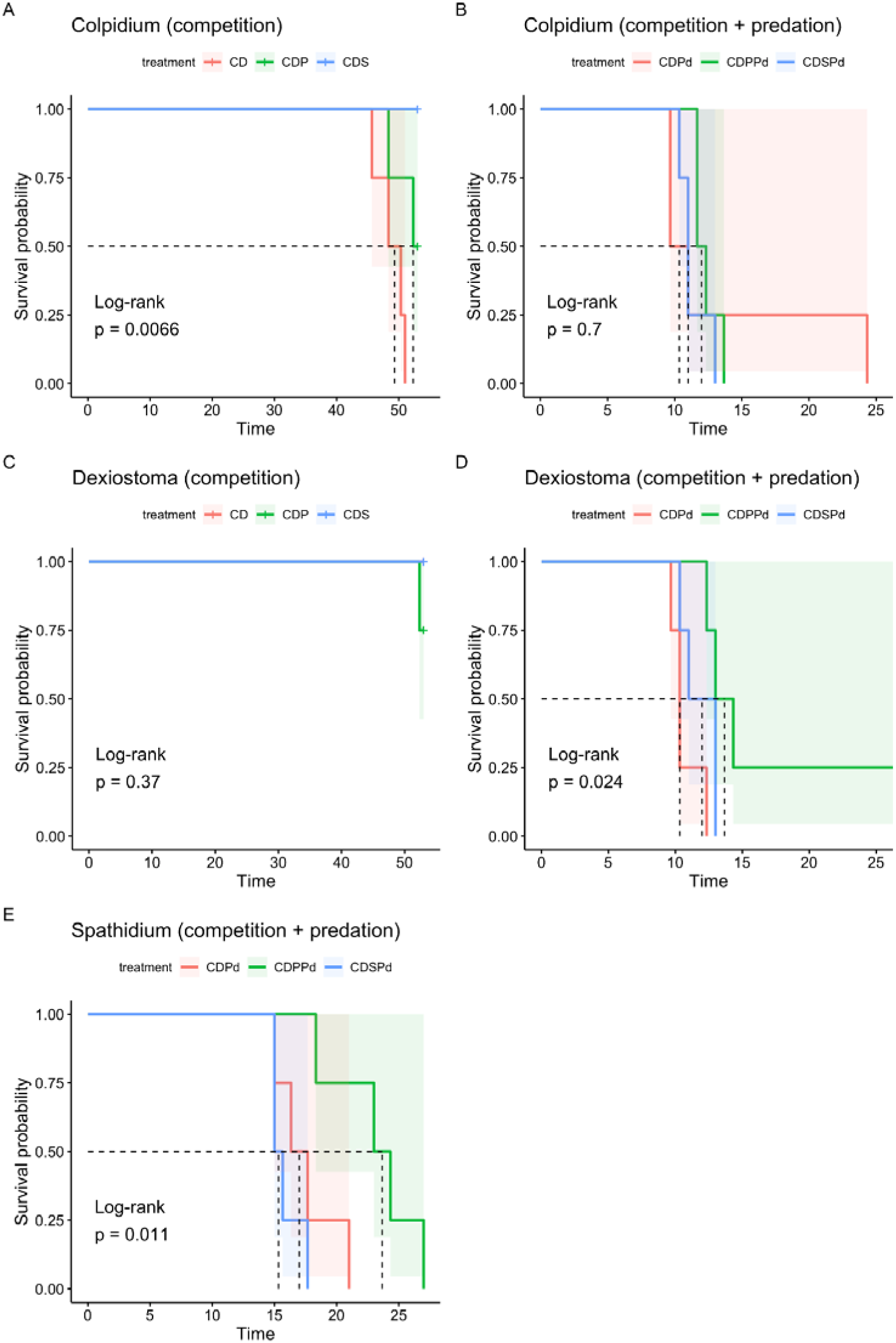
Survival curves (lines) of *Colpidium*, *Dexiostoma* and *Spathidium* as a function of community composition (CD = *Colpidium* & *Dexiostoma*, CDP = *Colpidium*, *Dexiostoma* & *Paramecium*, CDS = *Colpidium*, *Dexiostoma* & *Spirostomum*, CDPd = CD + predator, CDPPd = CDP + predator, CDSPd = CDS + predator). 95% confidence intervals are shown by shaded boxes. While for *Colpidium*, survival depends on additional species in the competitive communities (A) but not the communities with predation (B), the opposite is observed for Dexistoma (C-D). *Spathidium* survival depended on the prey community composition (E).

## Discussion

### Patterns of population abundance

Overall, species’ abundances and coexistence were influenced by community composition. *Colpidium* and *Dexiostoma* grew to lower densities when cultured with *Paramecium*, a sign of competition for shared resources. *Colpidium* abundance was not affected by the presence of *Spirostomum*. However, the mean abundance of *Dexiostoma* was lower in the presence of *Spirostomum*, suggesting competition between the two species, but to a lesser degree than with *Paramecium*. These patterns are in line with known phylogenetic relationships between these species, where *Colpidium* is most closely related to *Dexiostoma*, then to *Paramecium* and finally to *Spirostomum* (Violle et al., 2011). These relationships have been found to predict competitive exclusion due to the similarity in mouth size between species, which in turn defines the feeding niche of species (Violle et al., 2011).

Predation lowered the abundances of the two focal prey species, but this effect could be partially mitigated in the presence of additional species. For instance, *Colpidium* benefitted from the presence of *Paramecium* when predators were present, having a higher average abundance than would be expected if the effects of both predation and competition were additive. This might be explained by the interference of *Paramecium* with the predator, indirectly leading to a decrease in interaction strength between the prey and the predator. Ciliate predators show a preferred predator-prey size ratio of 8:1 (Hansen et al., 1994), explaining why some species may interfere with predator foraging. *Dexiostoma* similarly benefited from *Paramecium* when the predator was present, reaching on average higher abundance than when *Colpidium* and *Spirostomum* were present.

Surprisingly, both *Paramecium* and *Spirostomum* densities decreased in the presence of the predator, indicating that the predator had a negative effect on the two largest protist species. Furthermore, the abundance of the predator was greater when *Paramecium* was present compared to when it was cultured with only the focal species pair, or the focal pair and *Spirostomum*. It thus appears that *Paramecium* was consumed in addition to the two focal prey and that while *Spirostomum* was attacked, it was potentially too large to be consumed. This observation could be explained by the mode of prey capture employed by the predator *Spathidium*. The cell mouth (cytostome) of *Spathidium* is furnished with a rodlike tip (toxicyst) to 20 paralyze or kill other microorganisms for easier consumption (Fyda et al., 2005). But while the toxicyst make it possible to paralyze large prey, phagocytosis still requires that prey organisms can be engulfed.

### Evidence for higher order interactions

Comparing additive and interactive LV models revealed the significant effect of intraspecific HOIs on the population growth rate for both *Colpidium* and *Dexiostoma* in the two species communities (Figure 5A). While our experiment was not designed to test the underlying mechanisms that give rise to intraspecific HOIs, one plausible explanation could be the presence of an “information cascade”, wherein individual behaviors within the same species are regulated by the actions of others (Bikhchandani et al., 1992; Potts, 1984). Studies have shown that the behavior of neighboring fish influences the direction and speed of the school (Ioannou et al., 2011). If a fish senses a signal of danger and turns, it creates a pressure wave in the water, and other fish respond to this pressure wave by turning as well (Ioannou et al., 2011). Although ciliates do not have as elaborate sensory systems as fish, they are capable of sensing changes in local population density and adjusting their movement strategies accordingly (Fronhofer et al., 2015; Pennekamp et al., 2014). Since movement and foraging are often closely linked (Van Dyck & Baguette, 2005), it is possible that movement-related crowding effects could lead to the emergence of intraspecific HOIs.

**Figure 5:**
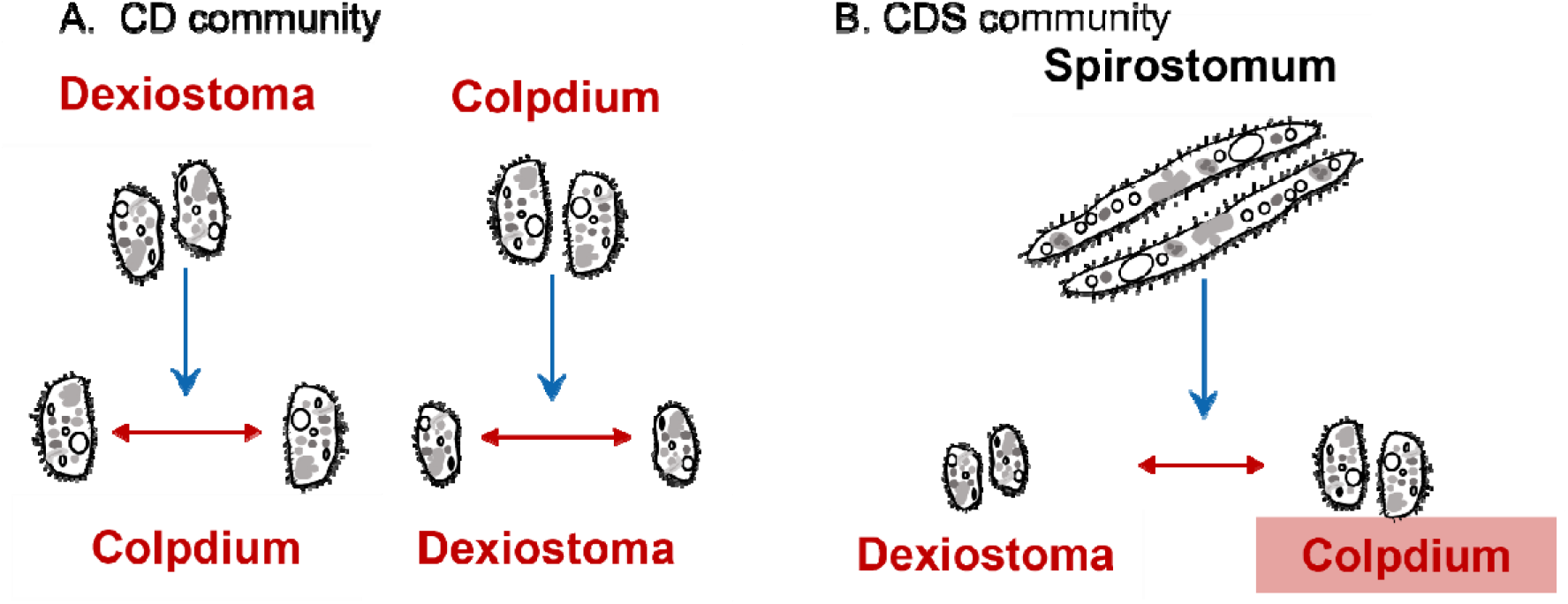
Higher-order interactions detected in our experiment. Direct pairwise interactions are shown as red arrows, HOIs are shown in blue. (A) In the two-species community of *Colpidium* and *Dexiostoma*, intraspecific HOIs were detected, indicating *Dexiostoma* modified the effect of *Colpidium* on itself while *Colpidium* modified the effect of *Dexiostoma* on itself. (B) In the community with *Colpidium*, *Dexiostoma*, and *Spirostomum*, an interspecific HOI was detected (The name of the affected species is marked with a light-colored box), suggesting *Spirostomum* modified the interaction between the two focal species.

Interspecific HOIs were only found on the population growth rate of *Colpidium* when *Dexiostoma* and *Spirostomum* were present (Figure 5B). *Spirostomum* most likely changed the effect of *Dexiostoma* on *Colpidium*, because both *Dexiostoma* and *Colpidium* strongly competed for the same resources, while the direct effect of *Spirostomum* on *Colpidium* was more limited. It is possible that there is an overlap in the size of the bacteria consumed by *Dexiostoma* and *Spirostomum*, while *Colpidium* uses a different size class. In such a case, *Dexiostoma* would compete with both species but would *Colpidium* only compete with *Dexiostoma.* This would also explain why we did not observe an interspecific HOI on the population growth rate of *Dexiostoma* when *Spirostomum* and *Colpidium* were present. Interestingly, there was no HOI of *Paramecium* on either *Dexiostoma* or *Colpidium*, although *Paramecium* was a stronger competitor than *Spirostomum*. Previous work has found strong evidence that interspecific HOIs can affect the coexistence and dynamics of aquatic microbial communities (Mickalide & Kuehn, 2019). Mickalide & Kuehn (2019) observed that *Escherichia coli* can invade cultures of the alga *Chlamydomonas reinhardtii* or the ciliate *Tetrahymena thermophila* but fails to invade a community where both species are present. The invasion resistance of the algae-ciliate community arises from an HOI caused by the algal inhibition of bacterial aggregation, which leaves bacteria vulnerable to predation.

The prevalence of intra-versus interspecific HOIs is noteworthy since most studies have so far focused on interspecific HOIs. Work on plant communities has shown that the prevalence of intra-or interspecific HOIs can vary across focal species and that neither type is generally more or less important (Mayfield & Stouffer, 2017). Since coexistence will be determined by the relative strengths of intra- and interspecific competition (Gibbs et al., 2022; Singh & Baruah, 2021), changes in intraspecific interaction strength alone can influence the persistence of communities. We therefore recommend that empirical studies investigating HOIs embrace a definition of HOIs that includes both intra- and interspecific effects (e.g., Mayfield & Stouffer, 2017).

### The nature of the detected HOIs

Some authors have argued that HOIs are not ecological processes in their own right but are instead emergent properties of phenomenological models (Letten & Stouffer, 2019). For example, a Lotka-Volterra competition model imposes a linear relationship between the density of one species and the growth of the other.

However, if this relationship is not linear in nature, the addition of HOI terms may improve the fit, but the model would not accurately describe the species’ interactions. Properly accounting for nonlinear density dependence is an important prerequisite to estimating interaction strength (Hart et al., 2018) and avoiding erroneous conclusions about the presence of HOIs when nonlinearity is present (Kleinhesselink et al., 2022; Letten & Stouffer, 2019). To investigate the potential for, and source of, nonlinear density dependence, we fit models that included either linear (i.e., additive LV) or non-linear (i.e., additive Ricker) competition coefficients, as well as non-linear intra- and interspecific HOI terms (i.e., the interactive models). We observed nonlinear density-dependence in all communities, as either the Ricker or the interactive LV model had the lowest AICc values. The observation that models which included HOIs had the lowest AICc suggests that there are processes resulting in non-linearity which the simple additive LV and the nonlinear Ricker model are missing. Therefore, we believe the HOIs detected are not an artifact of nonlinear density dependence, but true behavioural responses. A mechanistic model that allows for such non-linearity may provide further insights into the ecological processes driving community dynamics.

### Do HOIs affect abundance and community stability in competitive communities?

Persistence was different for the two focal prey species. *Dexiostoma* showed high persistence regardless of the additional species present, while *Colpidium* became more stable in the presence of a third species. The increased persistence of *Colpidium* when cultured with *Paramecium* was not driven by an HOI, since the additive Ricker model was best supported by the data in the CDP community. But the highest persistence of *Colpidium* was detected in the presence of *Dexiostoma* and *Spirostomum*, where the interactive model was best supported. This pattern is consistent with a stabilizing effect due to interspecific HOIs.

*Dexiostoma* showed a similar persistence across all combinations of competitors, despite a non-significant trend to lower persistence when *Paramecium* was present. *Dexiostoma* also showed variation in the presence of HOIs, but the pattern is not consistent with a stabilizing role of HOIs: when present with only *Colpidium*, intraspecific HOIs were observed and *Dexiostoma* persisted till the end of the experiment. When *Paramecium* or *Spirostomum* was added, HOIs were no longer detected but neither did the persistence change. This suggests that changes in HOIs do not always result in changes in the persistence of focal species. Direct interactions among species may be much more important.

The predator itself showed the highest persistence when feeding on communities that contained *Colpidium*, *Dexiostoma* and *Paramecium*. When feeding on the focal pair without competitors, it showed intermediate persistence times, while communities of the focal pair with *Spirostomum* persisted the least well. Hammill and colleagues (2015) found a similar effect for non-trophically interacting species in a food web: if species that do not interact trophically are present, prey persists for longer in diverse food webs. Since we could not quantify the interaction strength of the predator on *Colpidium* and *Dexiostoma* in the presence of *Paramecium* or *Spirostomum*, we cannot say whether the change in predator persistence was due to HOIs or simply the direct and indirect effects among the members of the food web. Quantifying the role of trophic interaction modifications with functional responses is an exciting area for future research (Terry et al., 2017).

### Limitations of our work

While our study provides insights into the occurrence and implications of higher order interactions in aquatic microcosms, the small sample size used in our experiment limits the generalizability of our findings. Detecting interactions usually requires larger sample sizes, unless deviations from the additive model are very large (Burgess et al., 2022). Due to the small sample size of our experiment, we may have missed important but weaker interactive effects. In addition, we were only able to examine the interaction of a single focal pair across a few community compositions. To completely disentangle the drivers of community persistence, we would need to measure all interaction strengths between species, which was logistically impossible in our study due to the long and high-quality time series. Since our study only investigated HOIs between a focal pair, it is possible that we missed additional interspecific HOIs between the focal pair and the predator. Future research with a larger sample size and a wider range of competitors and predators is warranted to further elucidate the complex nature of higher order interactions in ecological communities.

### Conclusions

The role of higher-order interactions in community dynamics remains a key issue in community ecology (Gibbs et al., 2022; Levine et al., 2017). Currently, very few empirical studies measure HOIs in communities of real species and study how higher-order interactions affect community stability. Our study demonstrates that HOIs can play a role in community stability, and provides some support for the stabilizing effects of HOIs on competitive communities shown in theoretical studies (Grilli et al., 2017). Our study further provides an example of how to identify HOIs by combining experiments with time-series analyses and hence pave the way for more studies that study the occurrence and consequences of HOIs in natural communities. Our study adds to the growing body of research that shows that species interactions can deviate from the pairwise expectations when additional species are present, and a new theory needs to explore the consequences for community coexistence and stability.

## Acknowledgments

We thank two reviewers and the associate editor for constructive feedback on the manuscript. Chenyu Shen, Kimberley Lemmen and Frank Pennekamp received financial support from the Swiss National Science Foundation (grant 310030_197811 awarded to FP). We thank Uriah Daugaard for inspiring the ciliate illustrations in Figure 1.

## Data and code accessibility

Data and code are available from the following github repository: <u>DOI:</u> <u>10.5281/zenodo.7896204</u>.

## Supplementary material

### S1. Preparation of bacterial cultures

One week before the start of the community experiment, we prepared the bacterial medium for prey cultures. We first added 3-4mL DMSZ medium into three 20mL Falcon tubes, then transferred *B. subtilis, B. brevis* and *S. fonticola* to these tubes using inoculation loops and incubated them at 37°C for 24 hours.

After one day of incubation, we added 100µl of *S. fonticola* and *B. subtilis* and 2ml of *B. brevis* medium (since *B. brevis* grows much slower than *S. fonticola* and *B. subtilis*) from the Falcon tubes to three previously autoclaved 1L glass bottles, each containing 500ml of ciliates pellet medium. We then incubated them at 20°C for two days to give the bacteria enough time to reach their carrying capacity. The prey culture medium was produced by mixing 33.3ml of each bacteria medium from the 1L glass bottles into an autoclaved 250 mL jar (i.e., the microcosm), yielding a total of 100ml of mixed bacterial culture medium. Two wheat seeds were added to each jar to provide a source of slow release nutrients that the bacteria could feed on, leading to stable populations of ciliates during our experiment (Altermatt et al., 2015). All the consumers were grown at 15° C prior to the experiments for three weeks to reach their carrying capacities.

### S2. Species classification

For species classification, we used the videos collected from the first sampling day of every week and randomly chose one video of each replicate for the six community compositions. By observing the morphology and movement patterns of the ciliates in the videos, we labelled 350 to 400 trajectories of each species as training data. We use a similar number of labelled trajectories of each species, which avoids low classification performance for rare classes (e.g. low abundance species) caused by imbalanced numbers of observations (Sommer & Gerlich, 2013). Although the trajectory filtering function from BEMOVI removed most of the background noise, the movements of some tiny, suspended impurities may remain as spurious trajectories within the training data. In order to prevent these trajectories from being mistakenly classified as other species, we selected 100 videos that we visually checked to contain no ciliates and labelled all detected trajectories as “noise”. Trajectories of class “noise” were then excluded from all subsequent analyses.

Twenty features extracted from an established classification pipeline (Pennekamp et al., 2017) were selected to train the classification algorithm to distinguish among classes based on information about body size and movement patterns (Table S1). As the phenotypes of ciliates may change over time (Pennekamp et al., 2017), the week number was added as the 21^st^ feature in the classification algorithm to enhance the accuracy of the prediction.

As the species included in each treatment were different, we built six customized models for classification that only contained the known species of each community. Prior to the experiment, we compared the random forest (RF) and Support Vector Machine (SVM) as our classifiers for their computation efficiency and reliability of results (Fernández-Delgado et al., 2014). We used the randomForest package (Liaw & Wiener, 2002) and e1071 packages that contain the *svm* function (Meyer et al., 2014.). We compared the performances of two classifiers by calculating the classification errors for all training data (Table S2). Since SVM gave us overall lower classification errors, we applied it to our six customized models and generated time series of change density change of species for each community.

### S3. Species abundances

**Table S1:**
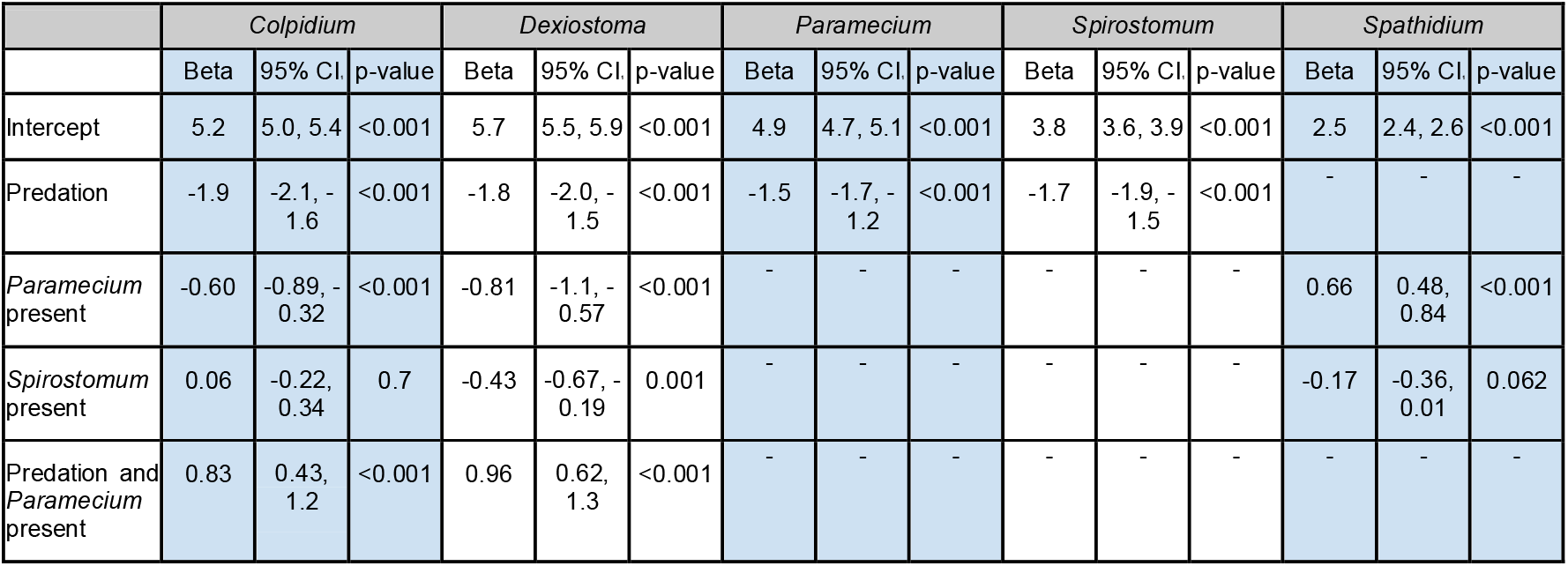
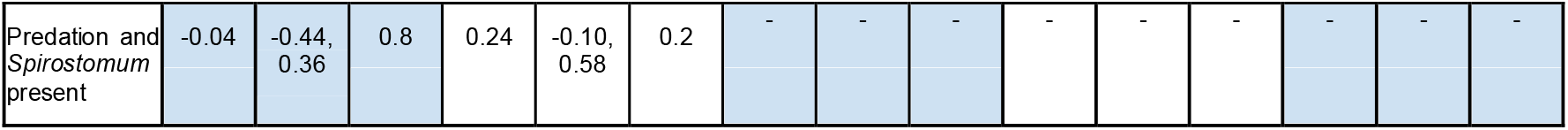
Effects of community composition on the log density of focal species. CI = confidence interval.

|  | *Colpidium* |  |  | *Dexiostoma* |  |  | *Paramecium* |  |  | *Spirostomum* |  |  | *Spathidium* |  |  |
| --- | --- | --- | --- | --- | --- | --- | --- | --- | --- | --- | --- | --- | --- | --- | --- |
|  | Beta | 95% CI | p-value | Beta | 95% CI | p-value | Beta | 95% CI | p-value | Beta | 95% CI | p-value | Beta | 95% CI | p-value |
| Intercept | 5.2 | 5.0, 5.4 | <0.001 | 5.7 | 5.5, 5.9 | <0.001 | 4.9 | 4.7, 5.1 | <0.001 | 3.8 | 3.6, 3.9 | <0.001 | 2.5 | 2.4, 2.6 | <0.001 |
| Predation | -1.9 | -2.1, -1.6 | <0.001 | -1.8 | -2.0, -1.5 | <0.001 | -1.5 | -1.7, -1.2 | <0.001 | -1.7 | -1.9, -1.5 | <0.001 | - | - | - |
| *Paramecium* present | -0.60 | -0.89, -0.32 | <0.001 | -0.81 | -1.1, -0.57 | <0.001 | - | - | - | - | - | - | 0.66 | 0.48, 0.84 | <0.001 |
| *Spirostomum* present | 0.06 | -0.22, 0.34 | 0.7 | -0.43 | -0.67, -0.19 | 0.001 | - | - | - | - | - | - | -0.17 | -0.36, 0.01 | 0.062 |
| Predation and *Paramecium* present | 0.83 | 0.43, 1.2 | <0.001 | 0.96 | 0.62, 1.3 | <0.001 | - | - | - | - | - | - | - | - | - |

|  |  |  |  |  |  |  |  |  |  |  |  |  |  |  |
| --- | --- | --- | --- | --- | --- | --- | --- | --- | --- | --- | --- | --- | --- | --- |
| Predation and *Spirostomum* present | -0.04 | -0.44, 0.36 | 0.8 | 0.24 | -0.10, 0.58 | 0.2 | - | - | - | - | - | - | - | - |

### S4: Model fitting to estimate interactions

The following section shows the coefficient estimates for the additive and interactive Lotka-Volterra and Ricker competition models.

**Figure S1:**
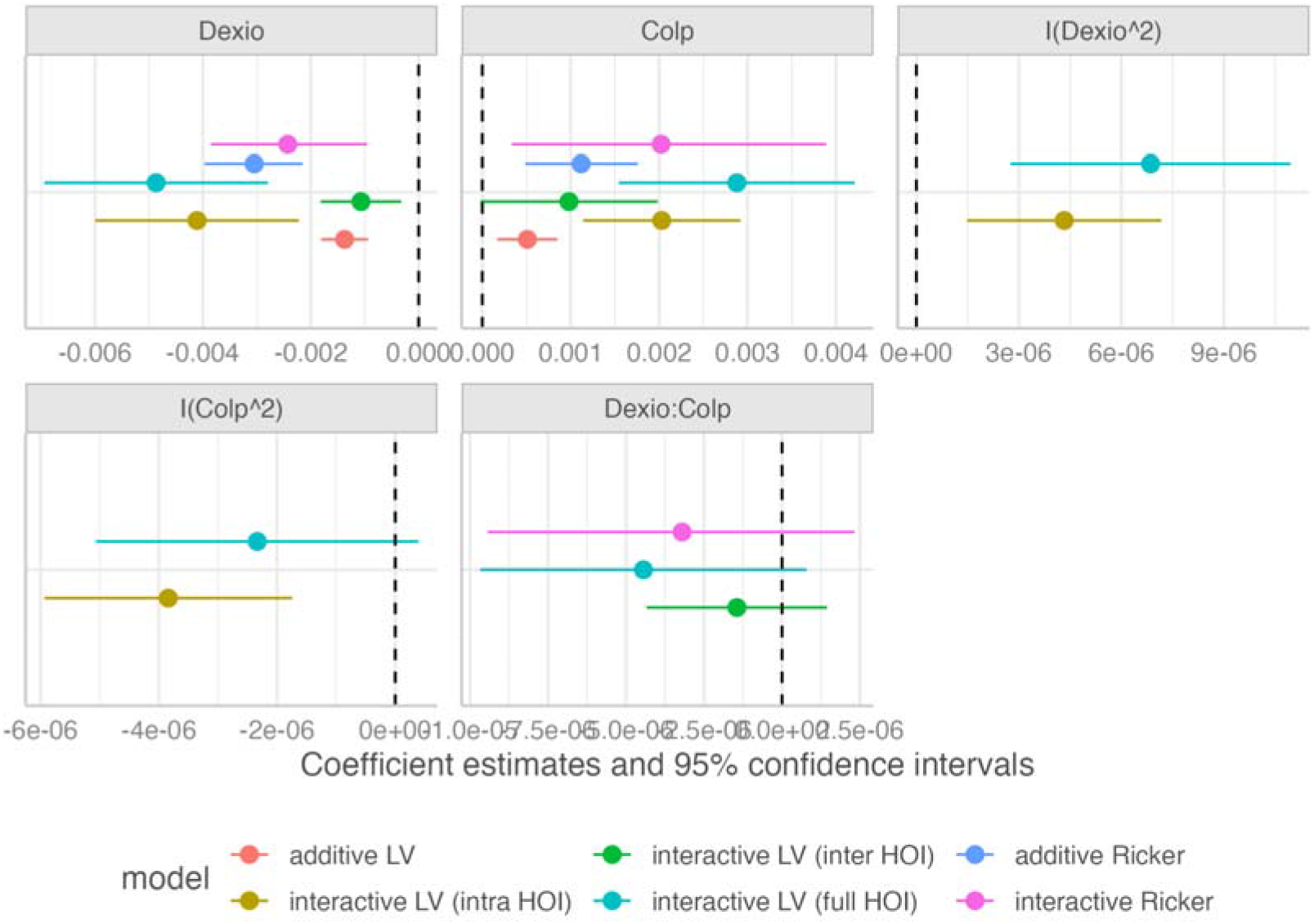
*Dexiostoma* population growth rate in CD community

**Figure S2:**
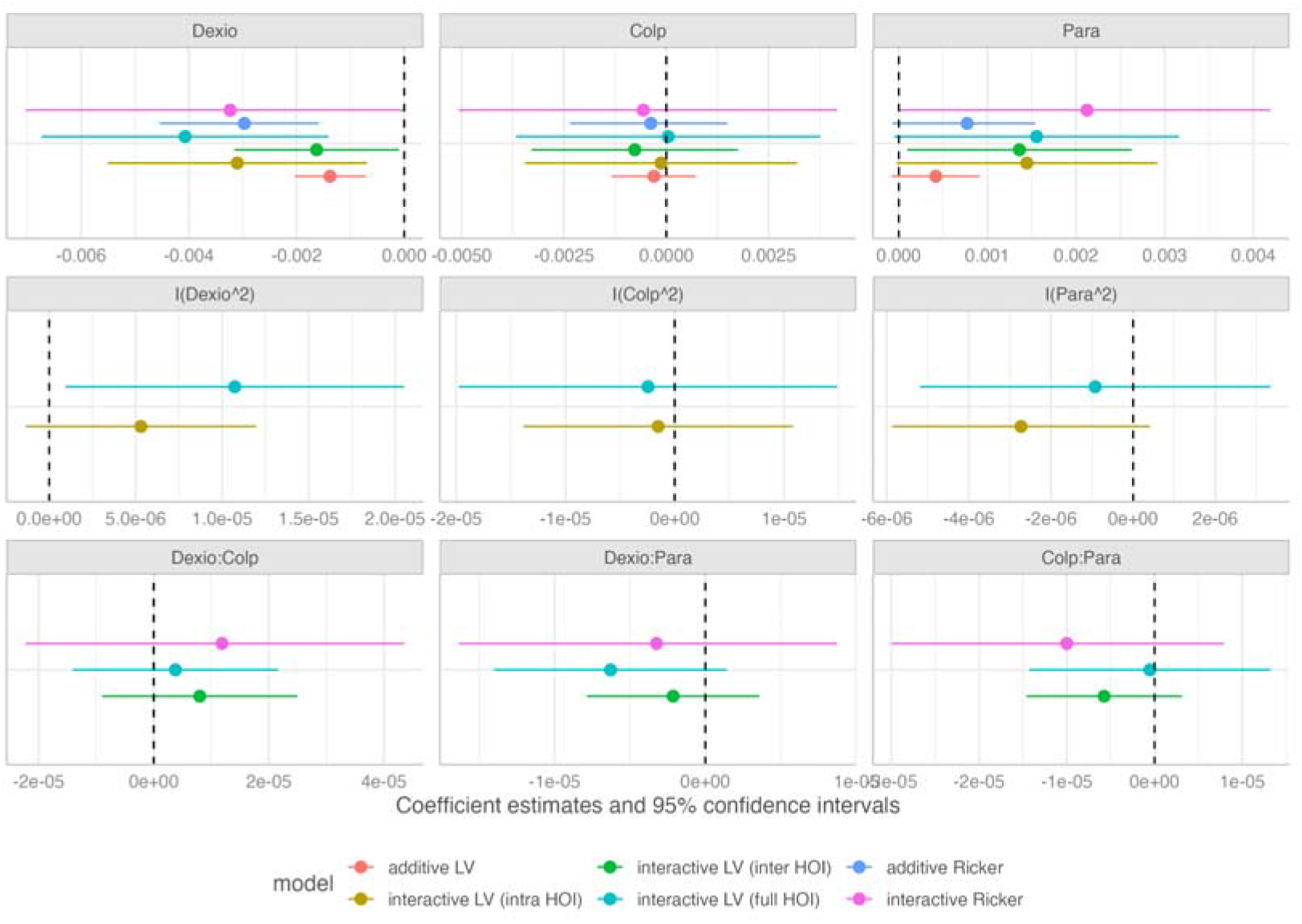
*Dexiostoma* population growth rate in CDP community

**Figure S3:**
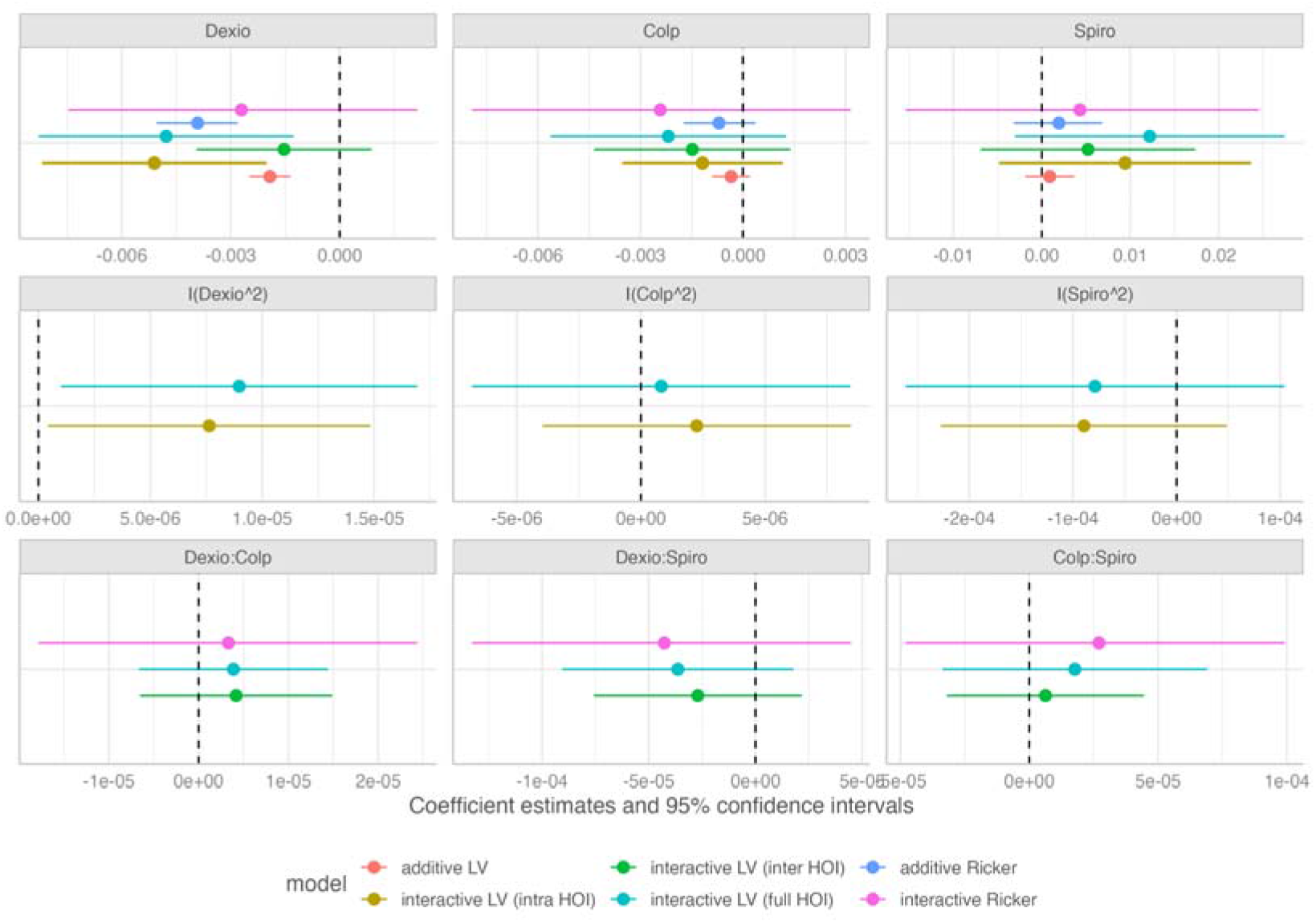
*Dexiostoma* population growth rate in CDS community

**Figure S4:**
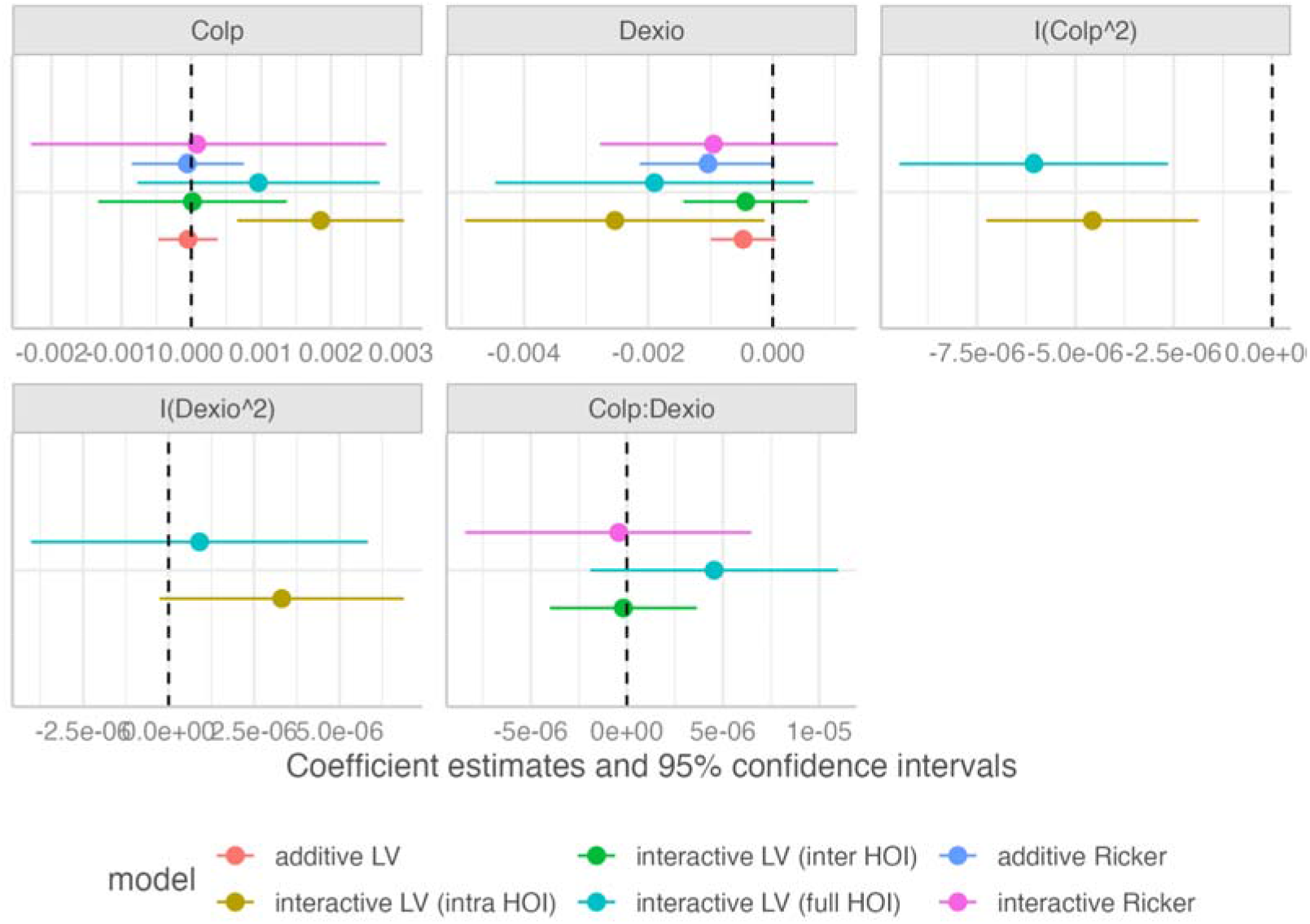
*Colpidium* population growth rate in CD community

**Figure S5:**
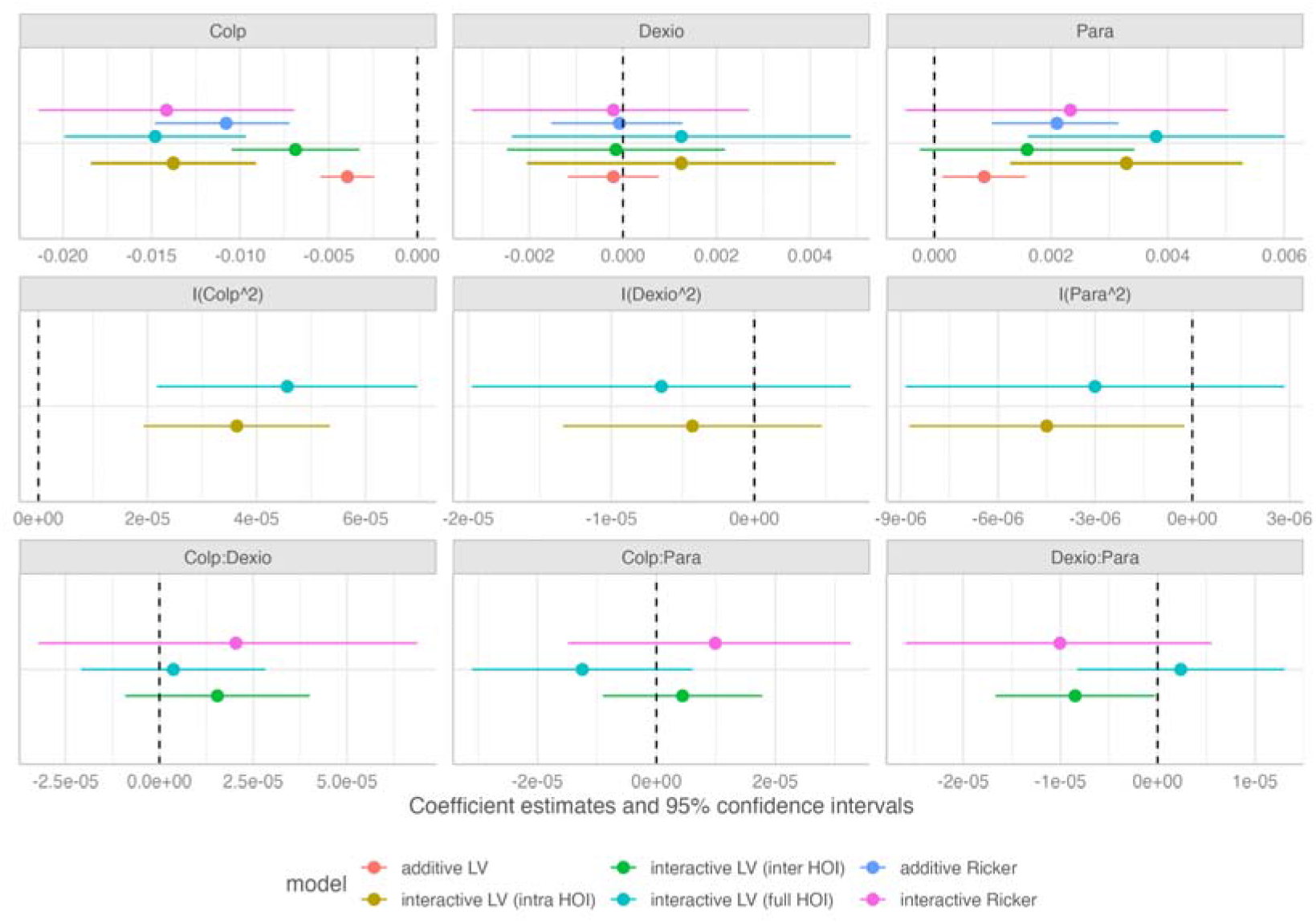
*Colpidium* population growth rate in CDP community

**Figure S6:**
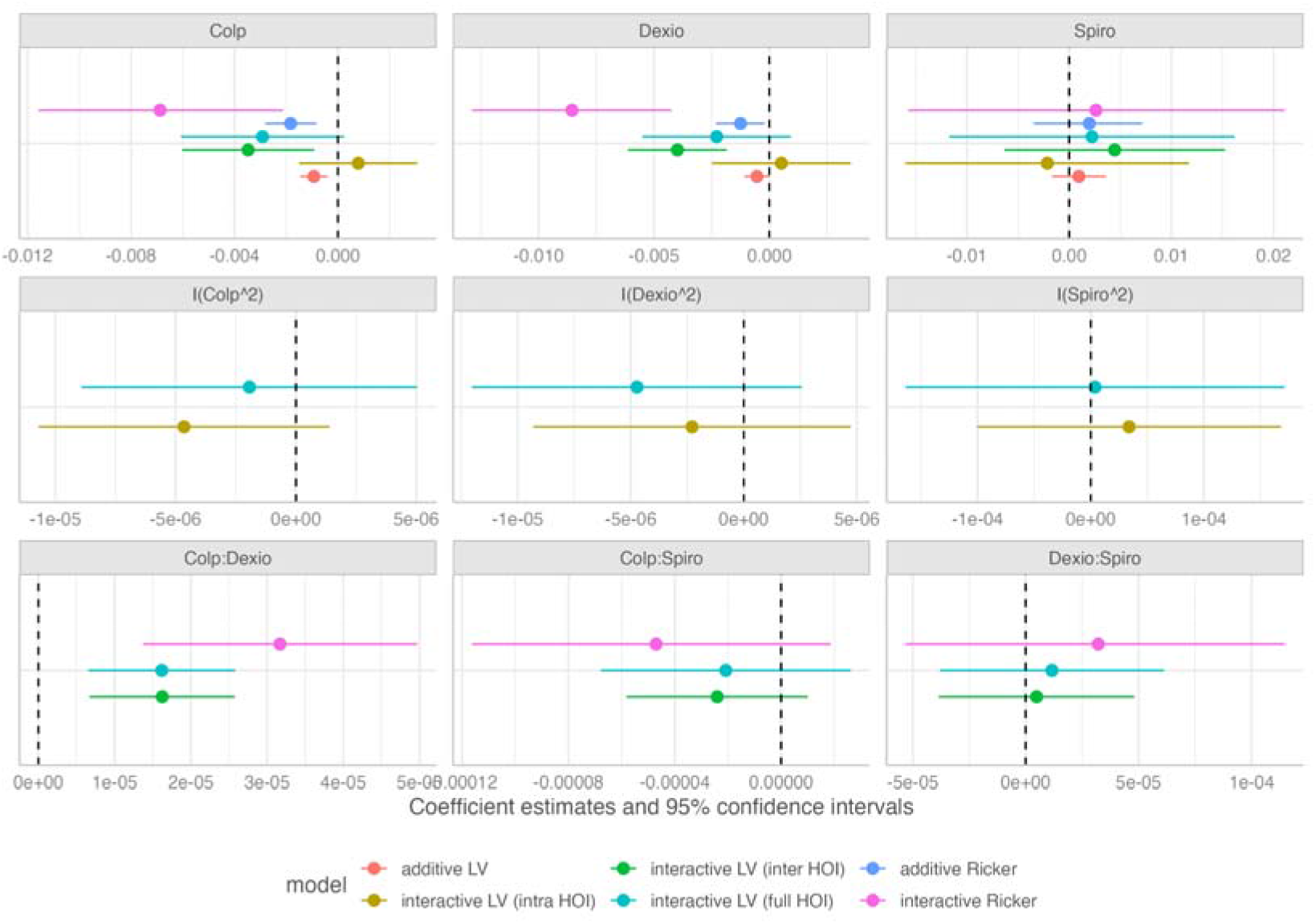
*Colpidium* population growth rate in CDS community

**Figure S7:**
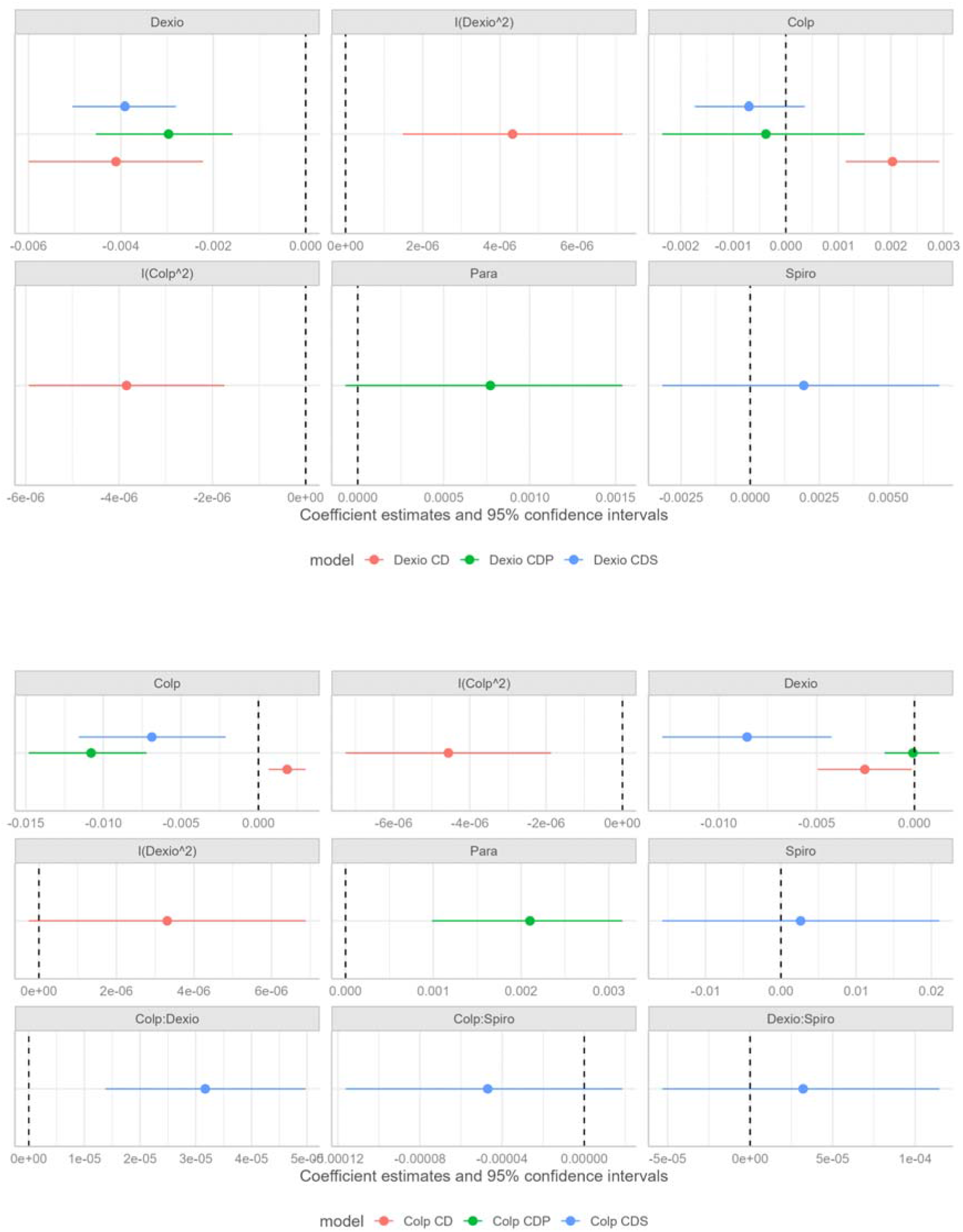
Coefficient plots for the best supported models for *Dexiostoma* (upper panel) and *Colpidium* (lower panel) across community compositions. Each panel shows the intra-or interspecific interaction terms as well as the interaction coefficients.

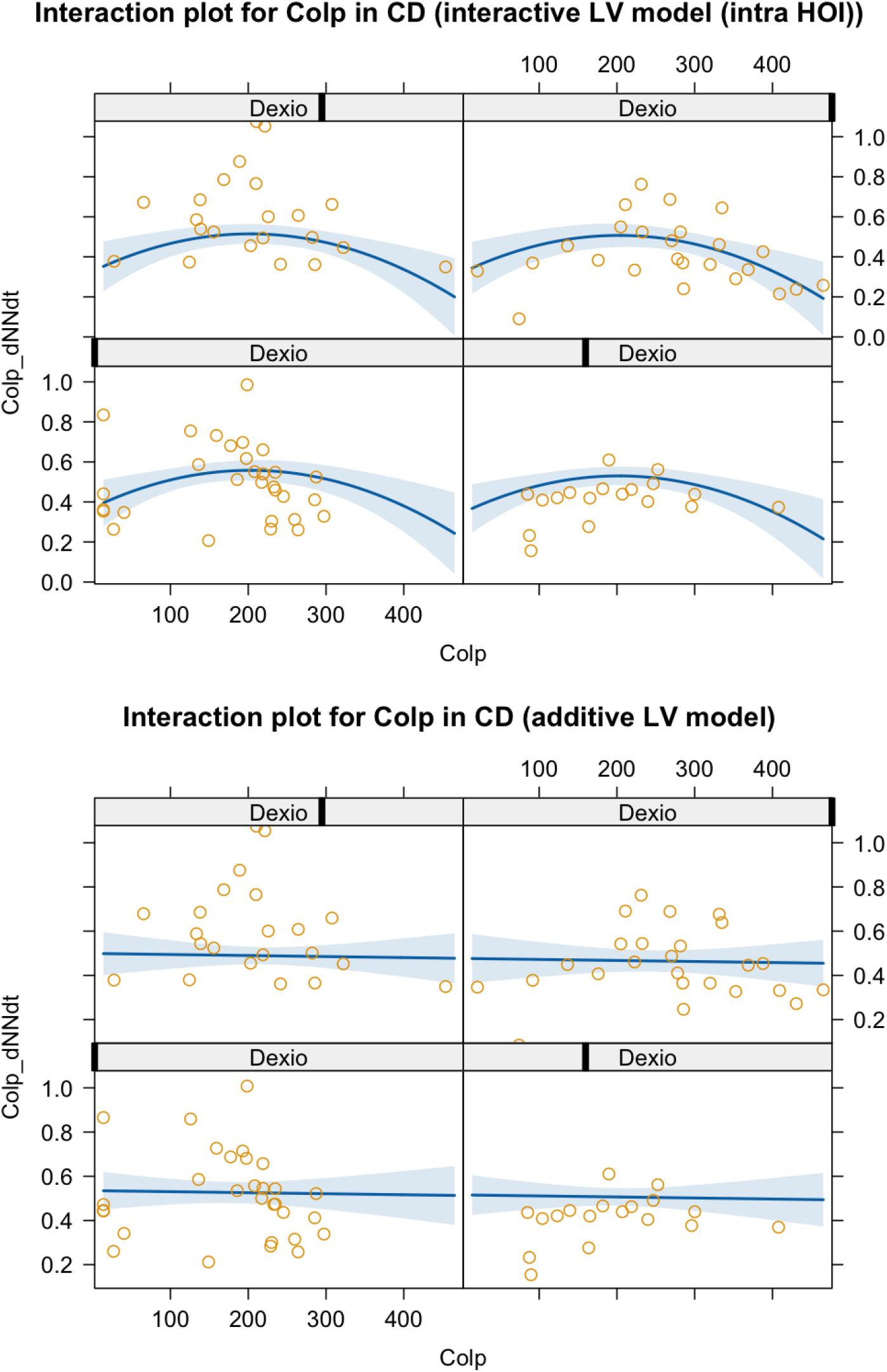
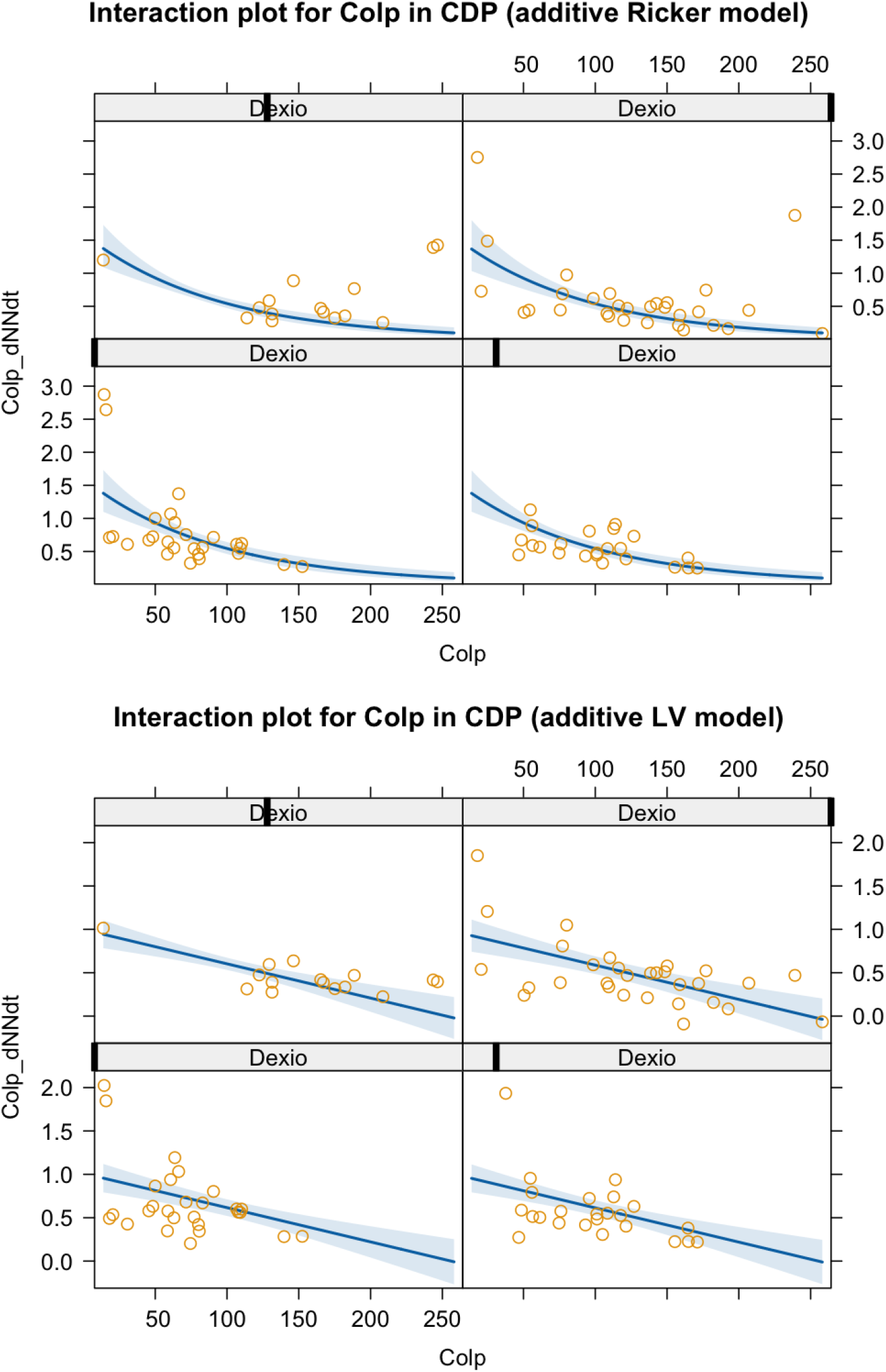
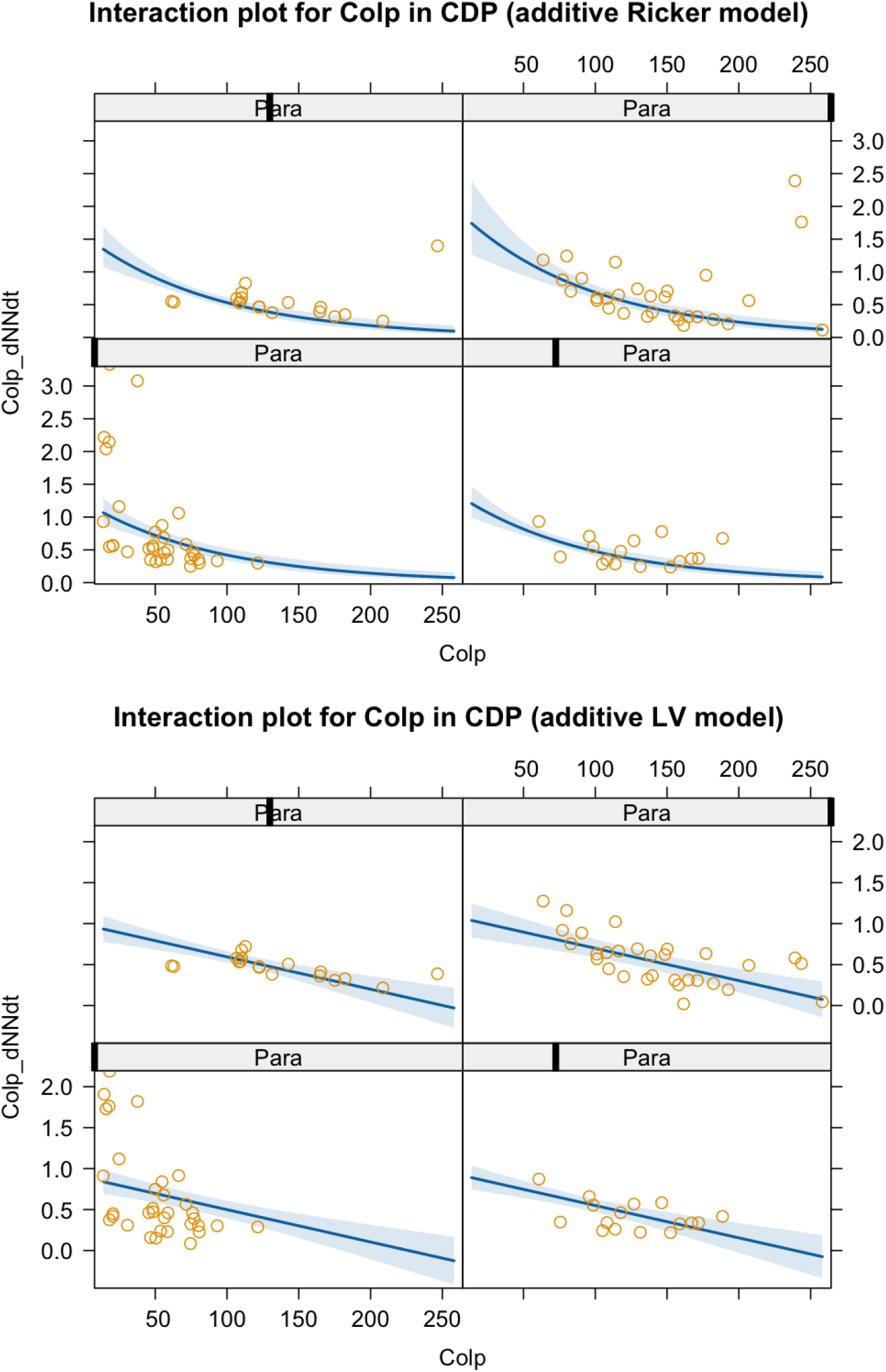
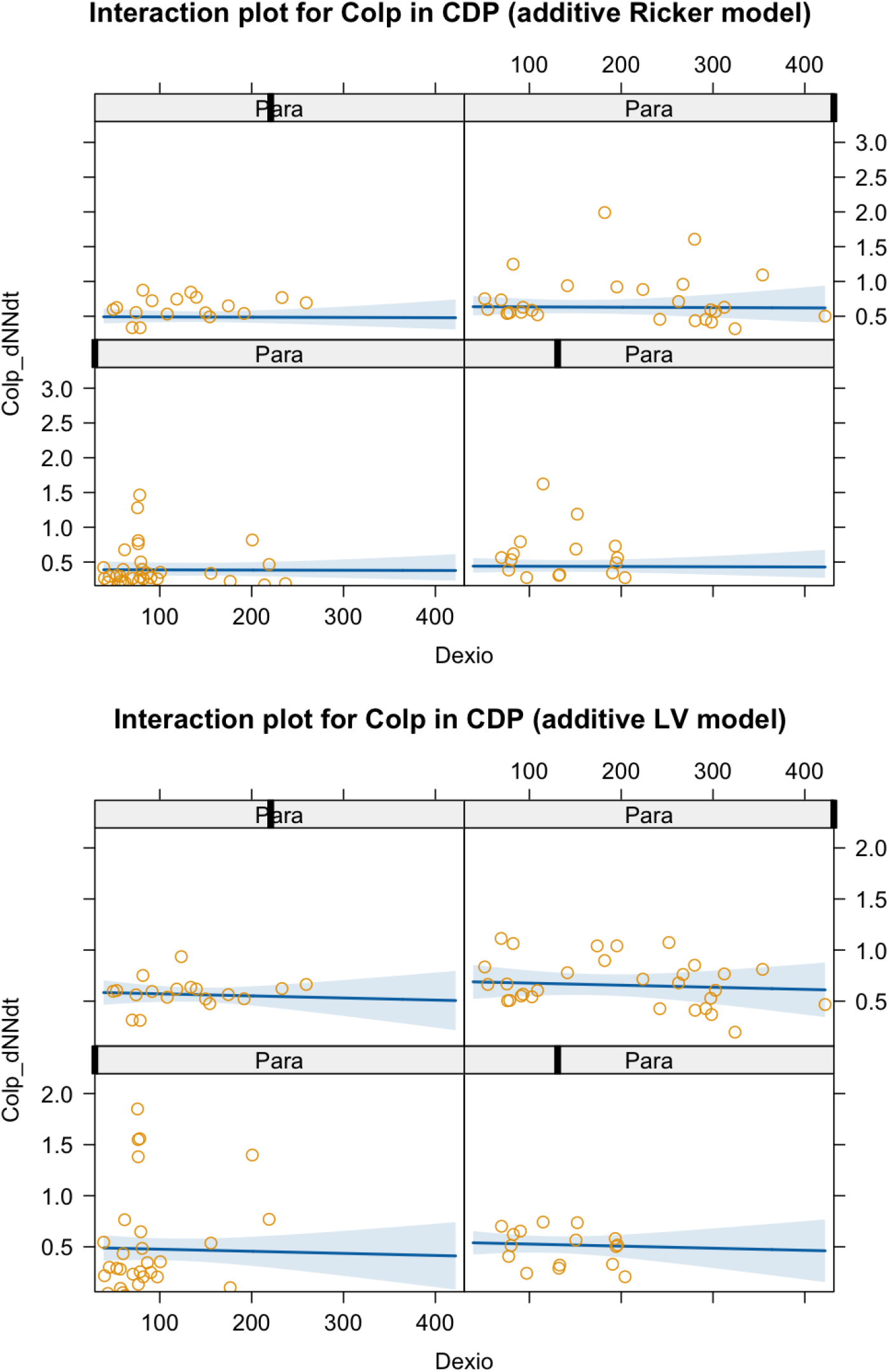
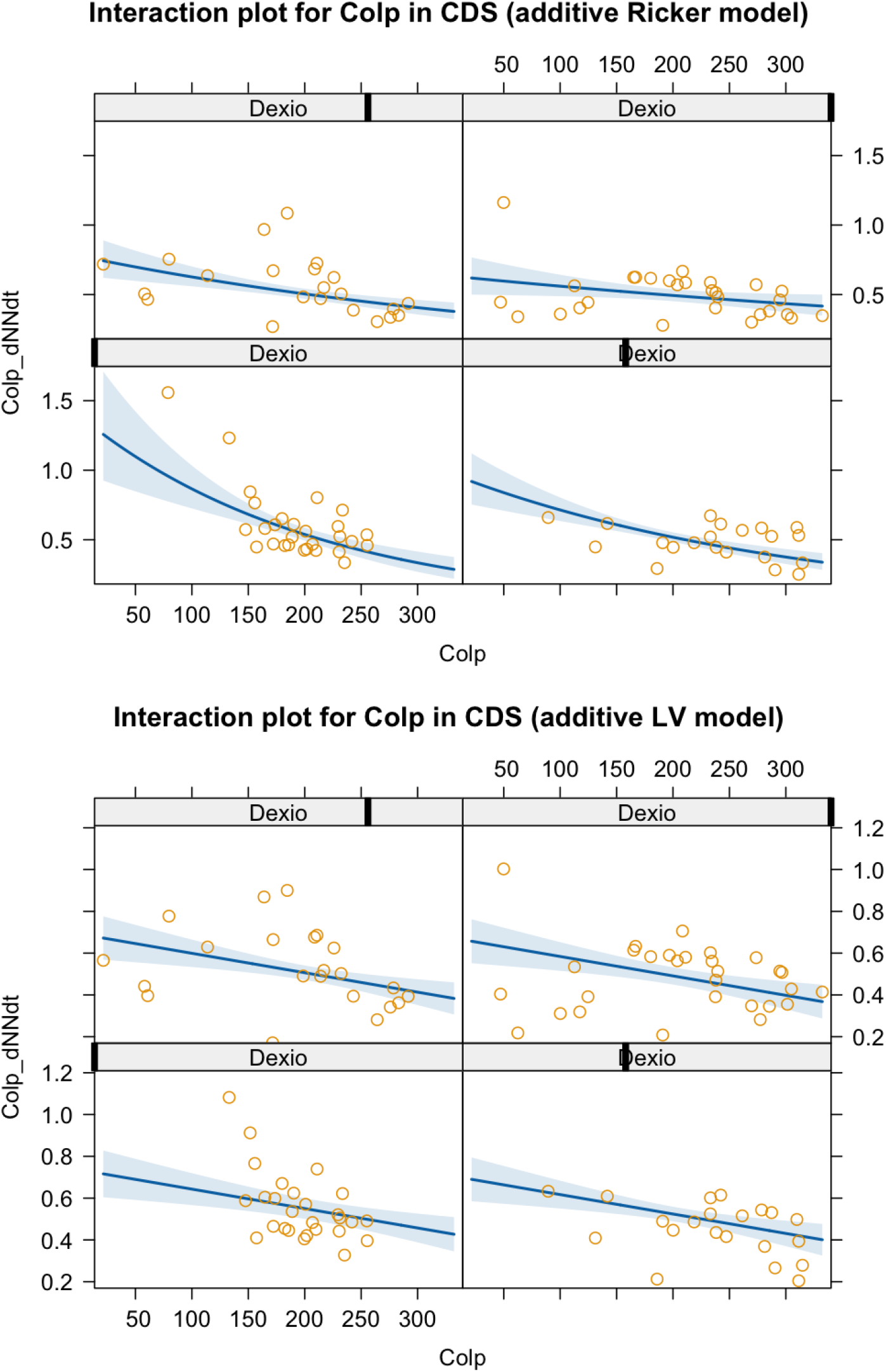
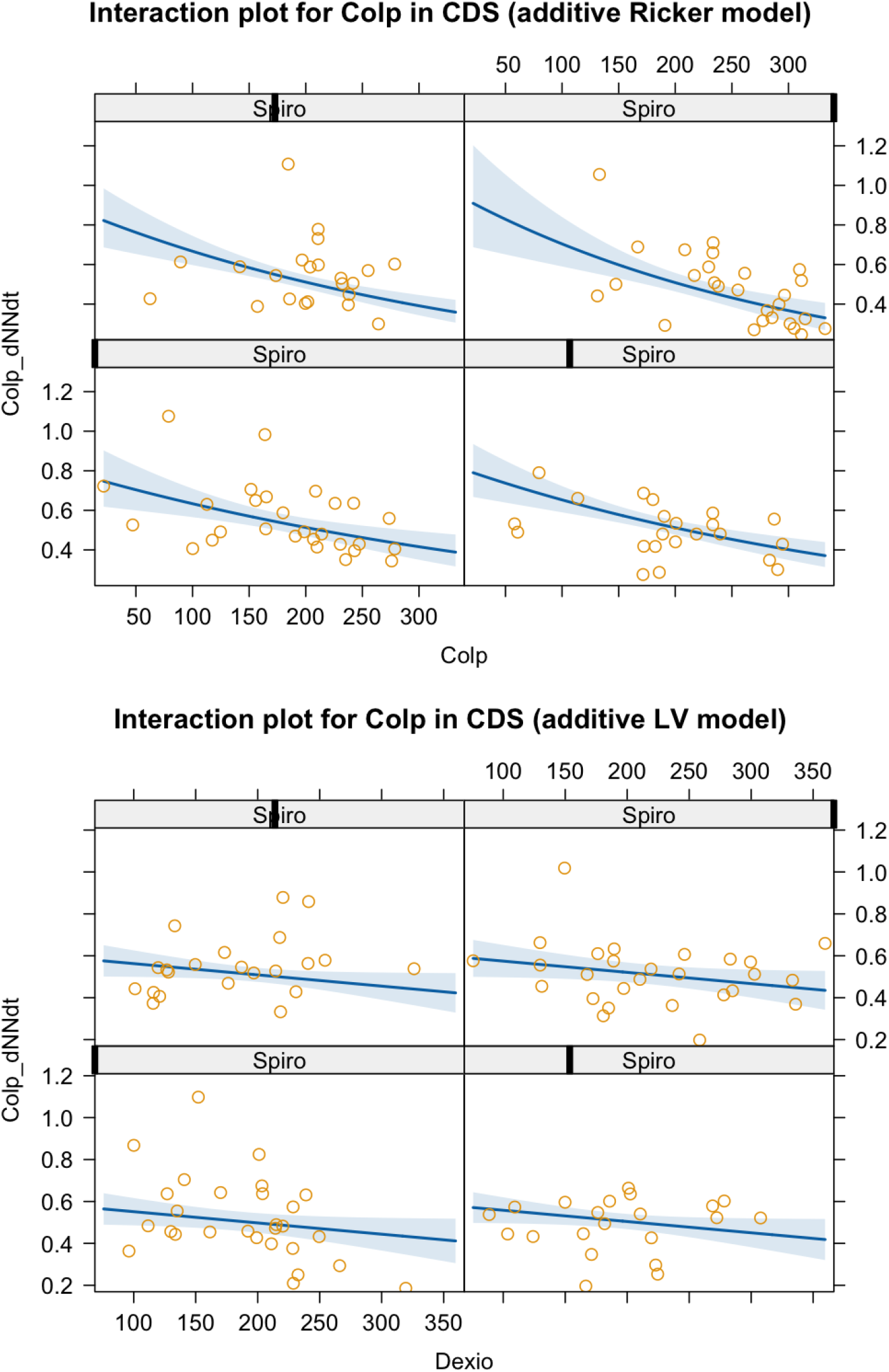
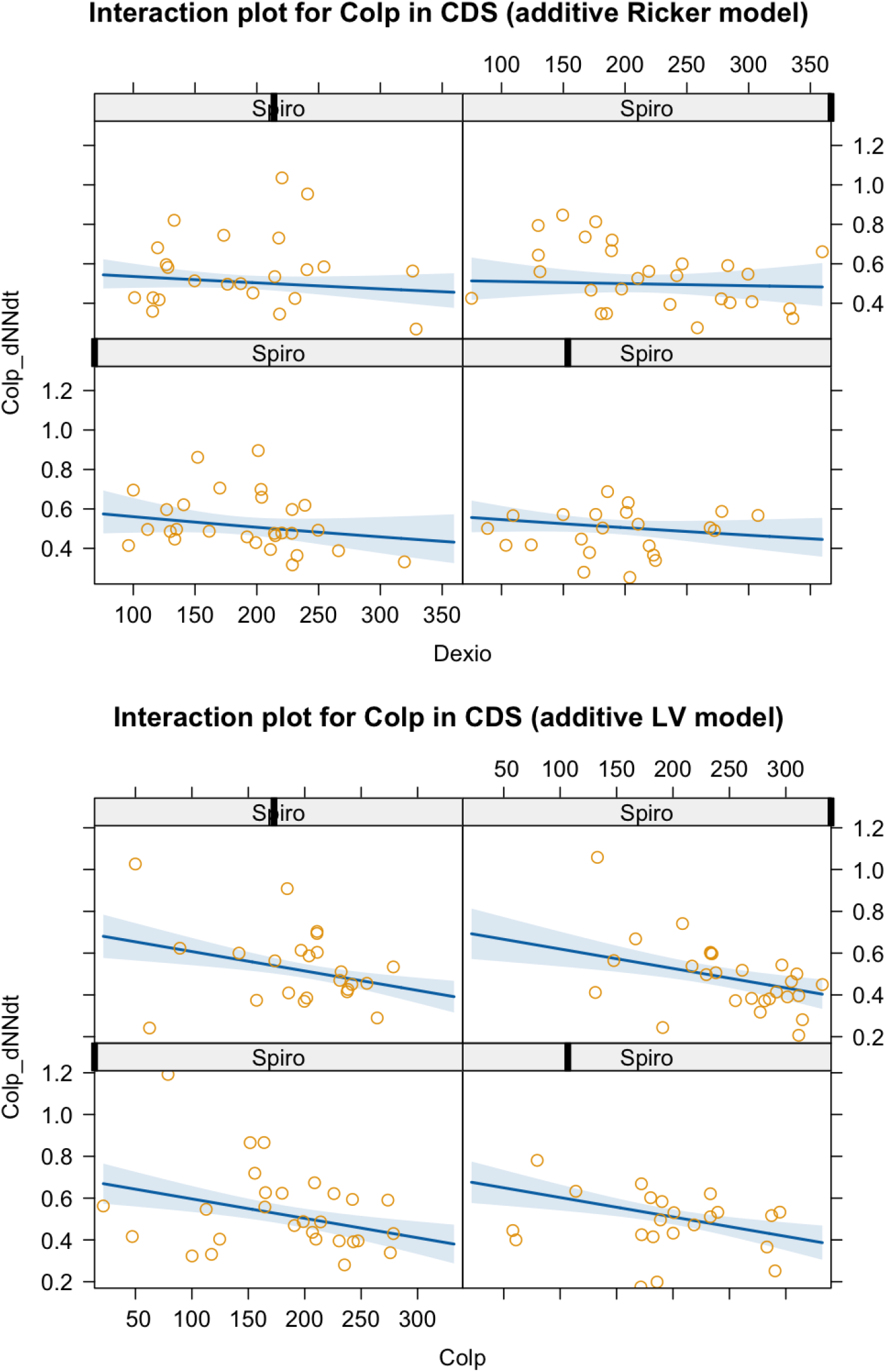
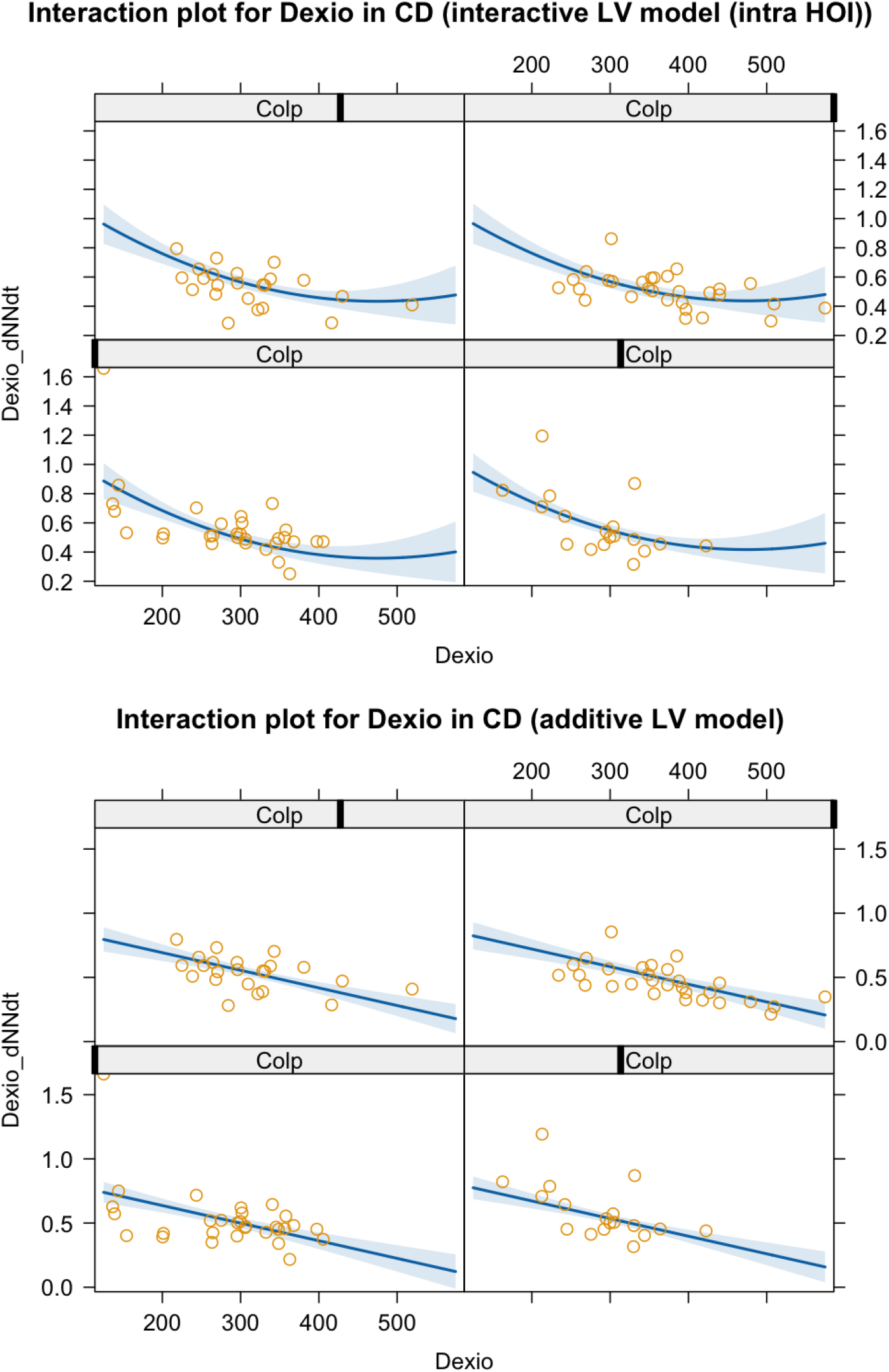
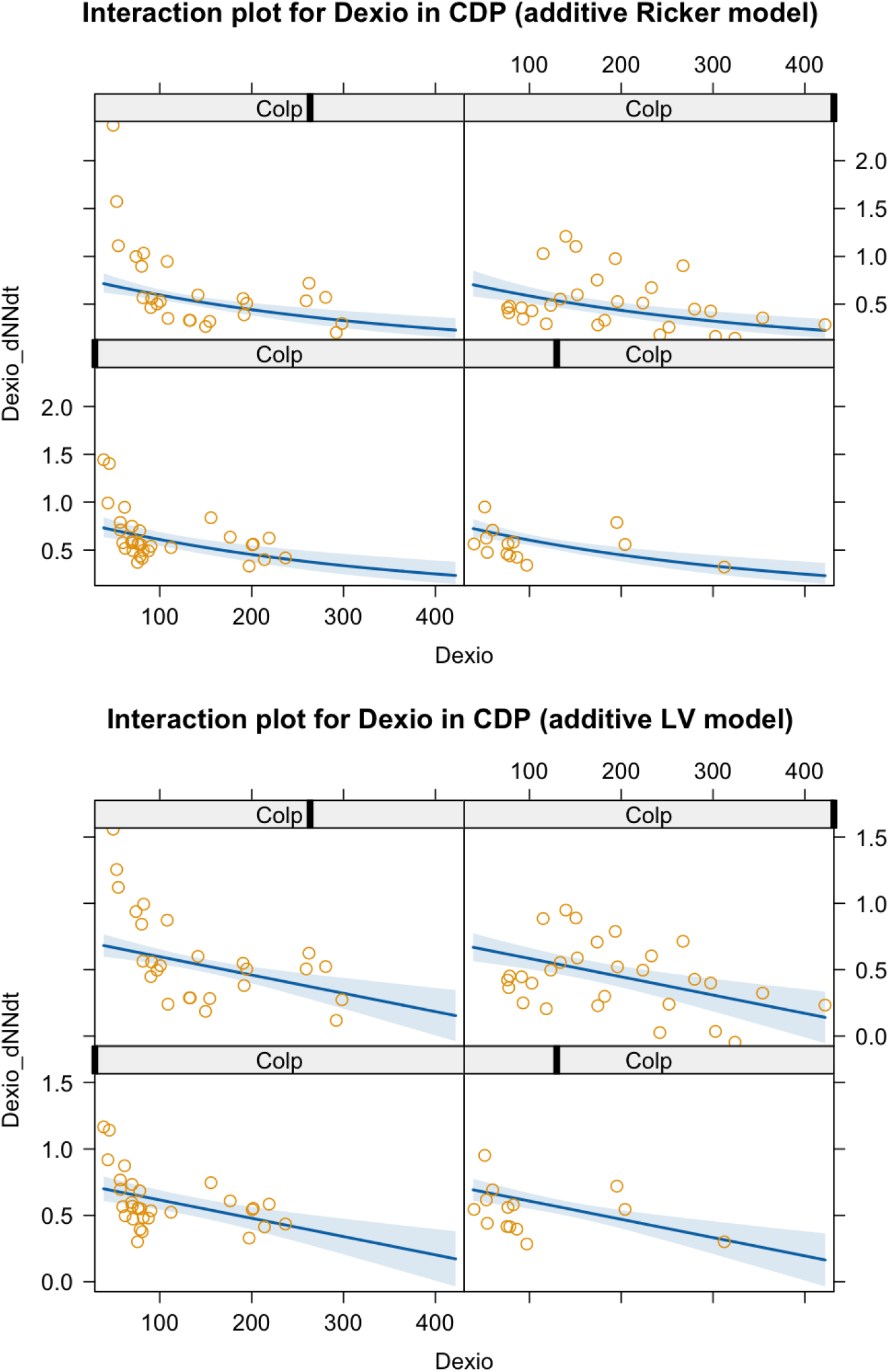
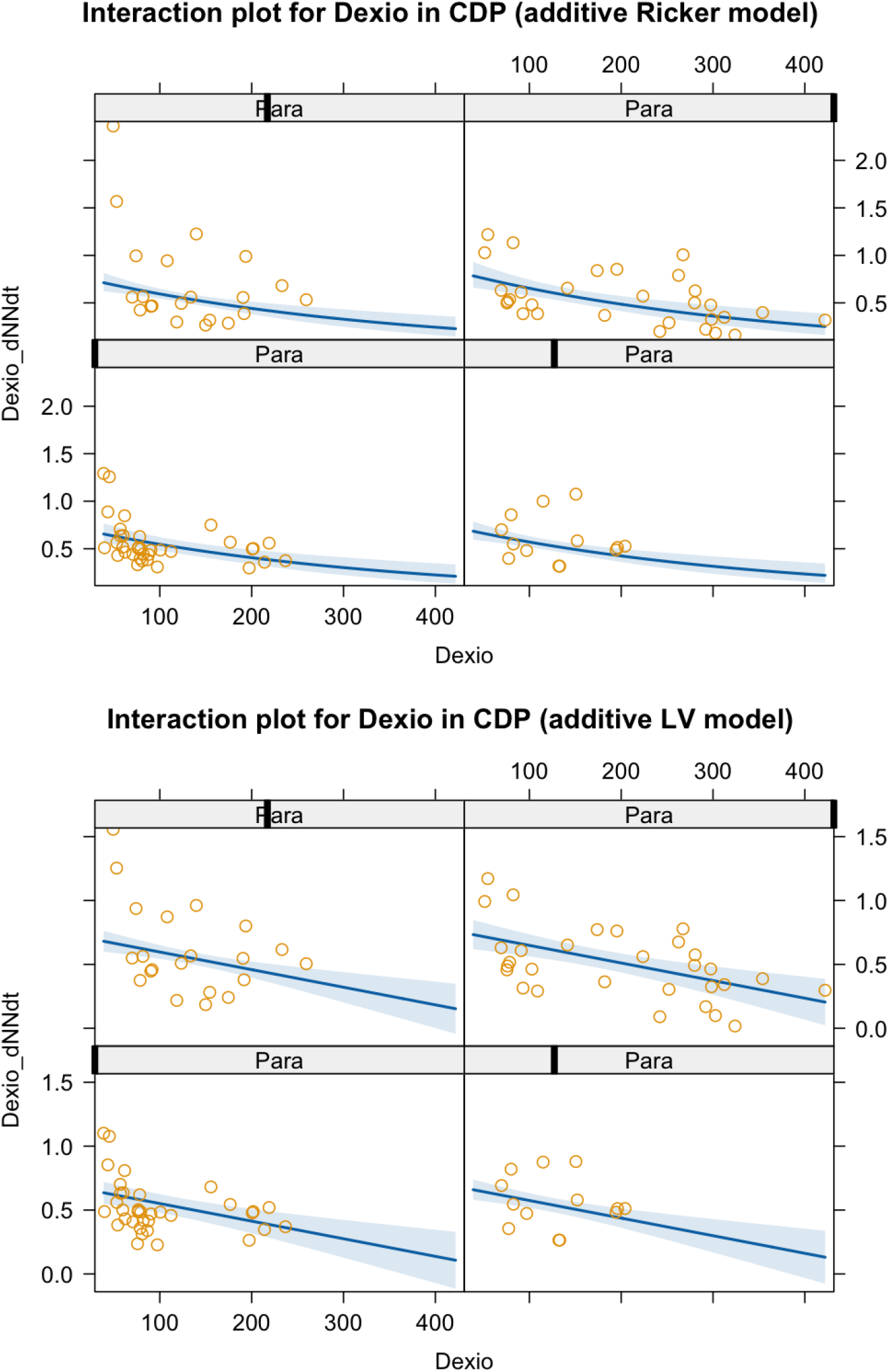
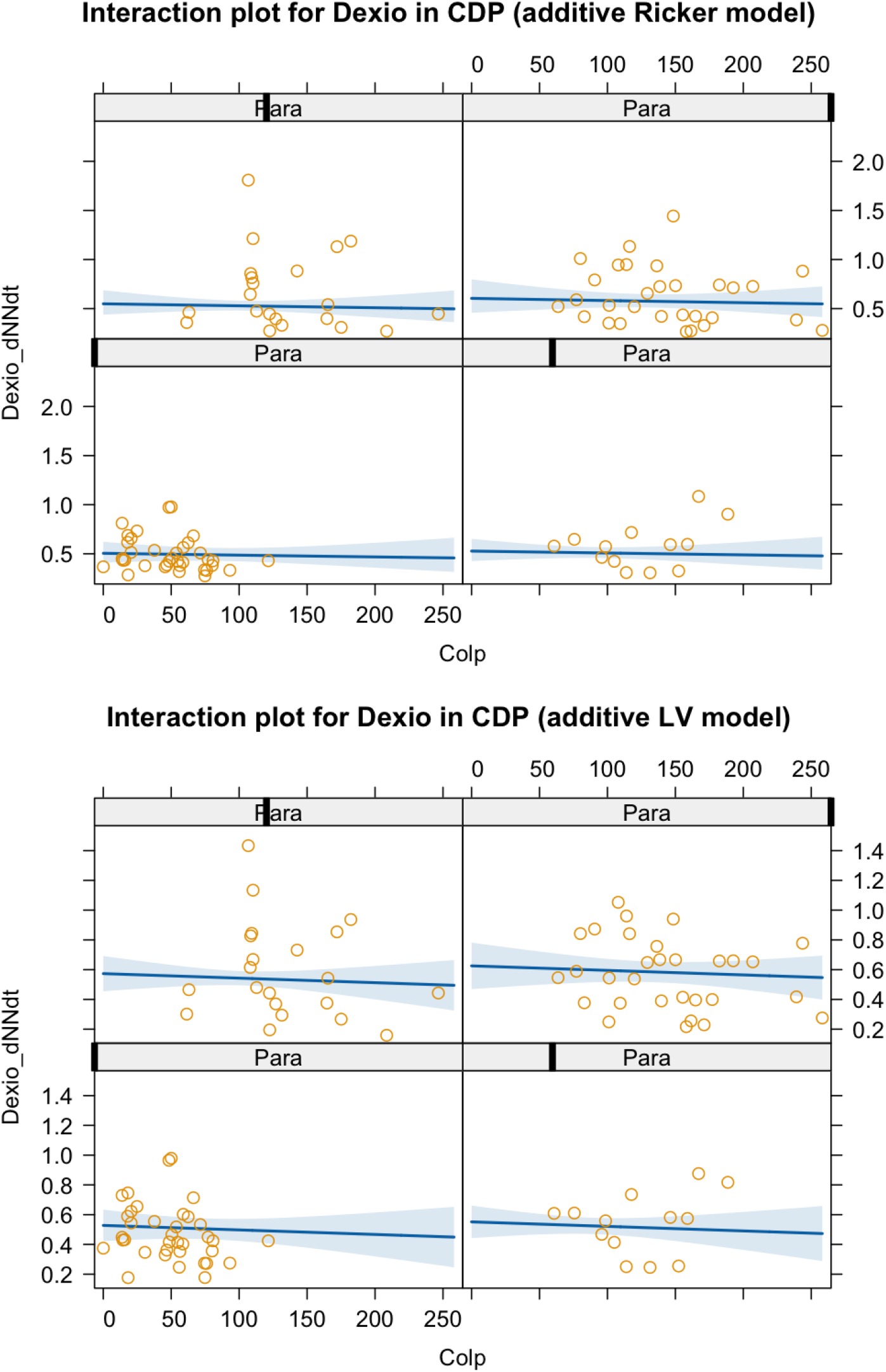
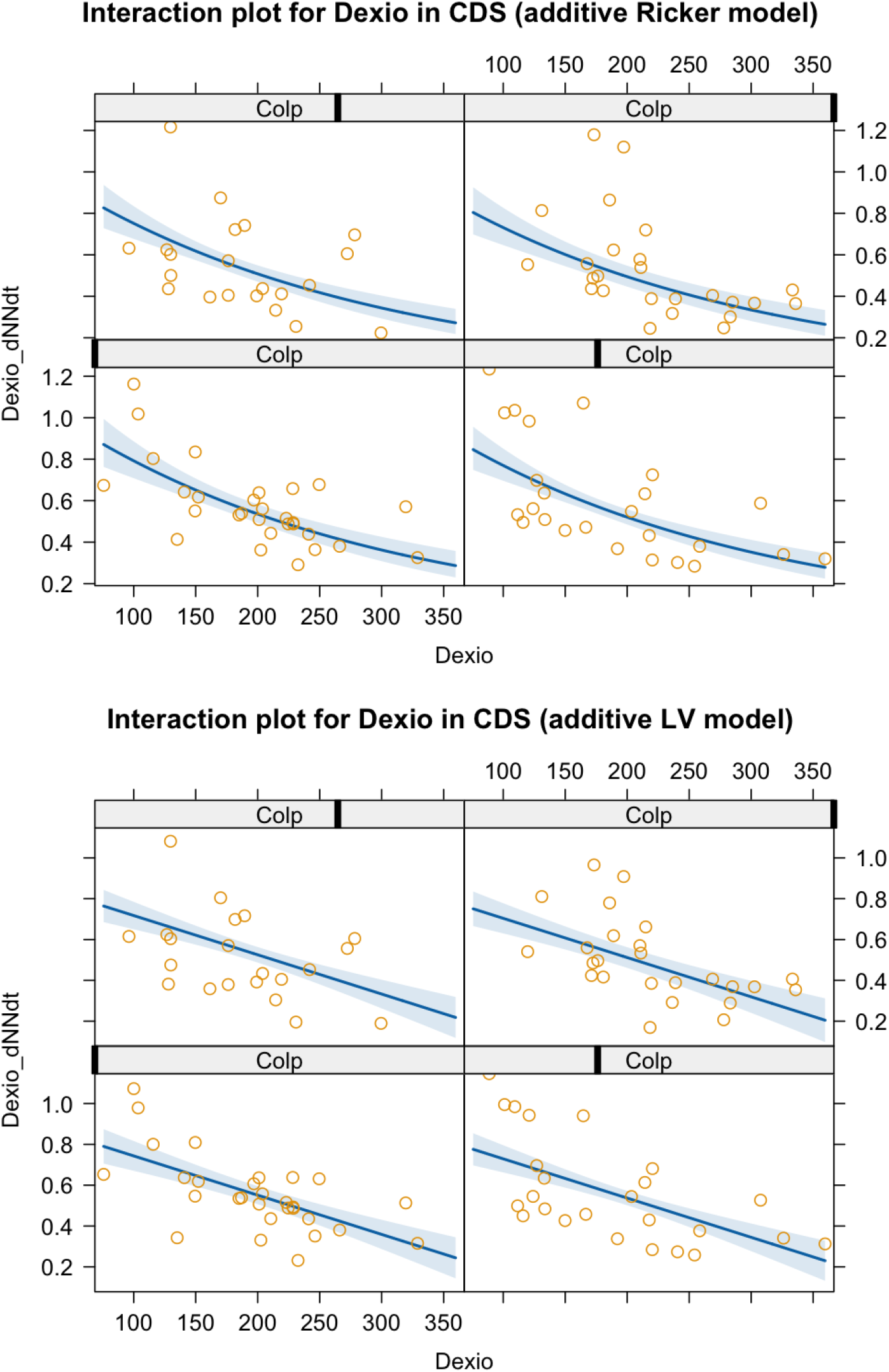
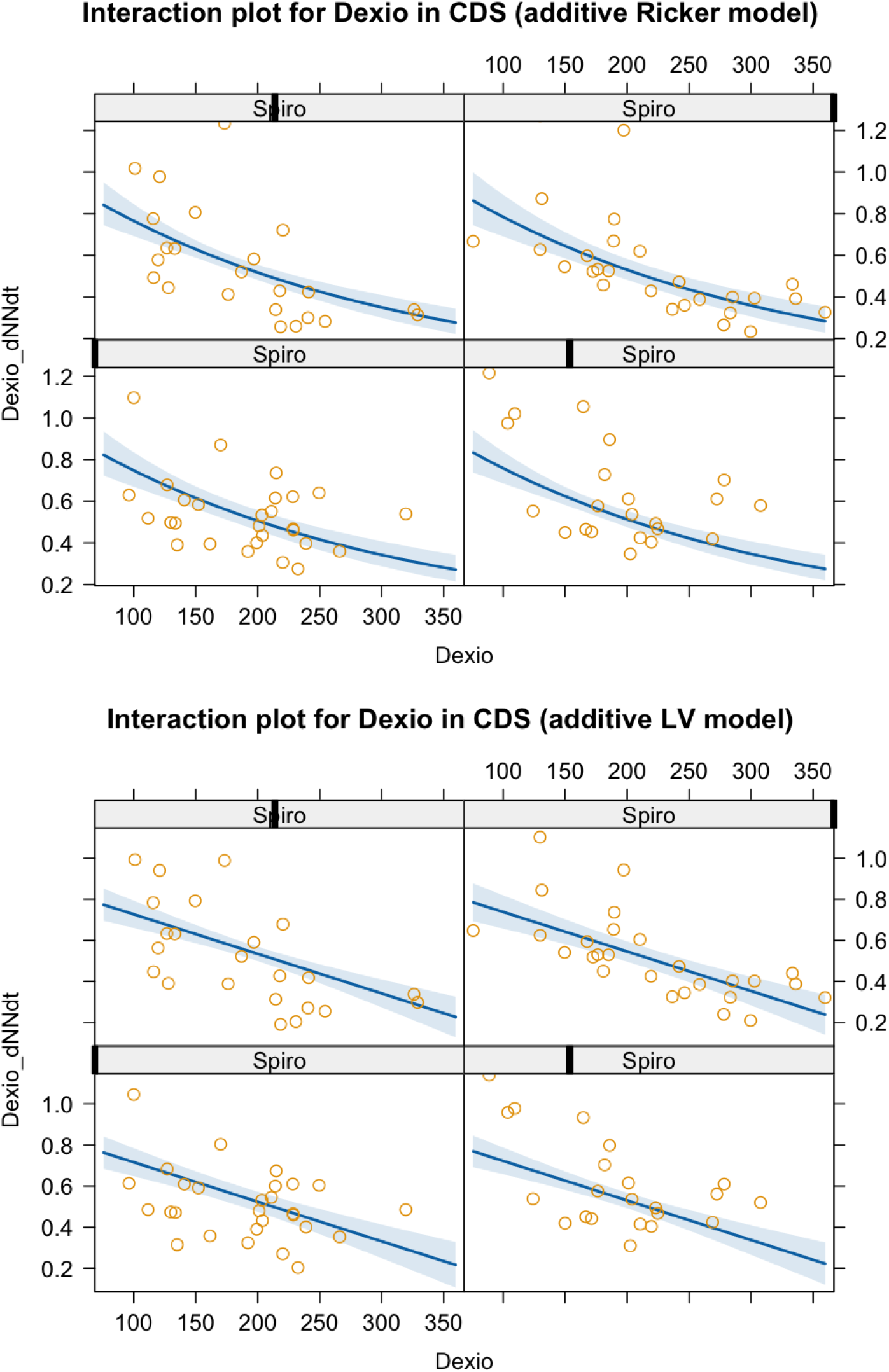
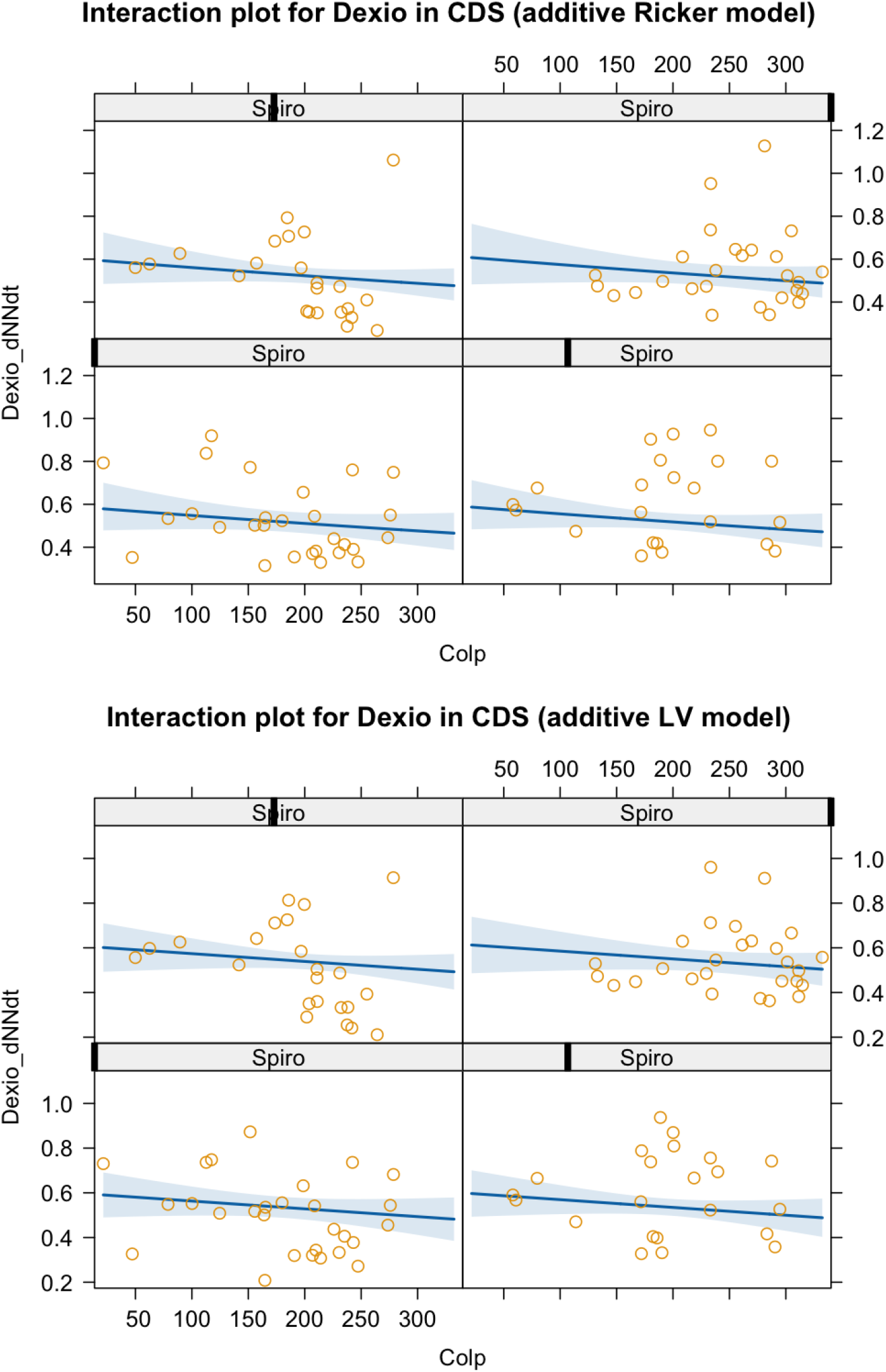

### S5. Model fit

We show the fit of the most parsimonious model compared to the additive LV model first for Colpidium, then for Dexiostoma, across all community compositions. The Y shows the population growth rate, whereas the x-axis shows the effects of intra-or interspecific density. Changes in model fit across panels show intra-or interspecific higher order interactions.

